# Tuned geometries of hippocampal representations meet the demands of social memory

**DOI:** 10.1101/2022.01.24.477361

**Authors:** Lara M. Boyle, Lorenzo Posani, Sarah Irfan, Steven A. Siegelbaum, Stefano Fusi

## Abstract

Social recognition consists of multiple memory processes, including the detection of familiarity – the ability to rapidly distinguish familiar from novel individuals – and recollection – the effortful recall of where a social episode occurred and who was present. At present, the neural mechanisms for these different social memory processes remain unknown. Here, we investigate the population code for novel and familiar individuals in mice using calcium imaging of neural activity in a region crucial for social memory, the dorsal CA2 area of the hippocampus. We report that familiarity changes CA2 representations of social encounters to meet the different demands of social memory. While novel individuals are represented in a low-dimensional geometry that allows for rapid generalization, familiar individuals are represented in a higher-dimensional geometry that supports high-capacity memory storage. The magnitude of the change in dimensionality of CA2 representations for a given individual predicts the performance of that individual in a social recognition memory test, suggesting a direct relationship between the representational geometry and memory-guided behavior. Finally, we show that familiarity is encoded as an abstract variable with neural responses generalizing across different identities and spatial locations. Thus, through the tuning of the geometry of structured neural activity, CA2 is able to meet the complex demands of multiple social memory processes.

## Introduction

Social memory, an animal’s ability to recognize and remember experiences with other individuals of its own species (conspecifics), consists of several discrete psychological processes. As illustrated by the classic example of “the butcher on the bus”^1^, these include the ability to rapidly detect whether an individual is novel or familiar (“I know that person but from where?”) and the more effortful recollection of an individual’s specific identity and the associated set of past experiences with that individual (“Ah, she’s my butcher, I bought food from her last Tuesday”). These processes have conflicting demands and requirements. Familiarity must readily generalize to allow the detection of whether an individual is novel or familiar across different contexts. In contrast, social episodes consist of complex multi-dimensional experiences with a particular individual at specific places and times and require distinct representations of individual identity even when the degree of familiarity is the same. How does the brain manage these conflicting memory requirements of familiarity detection, representation of social identity, and storage of social episodic memories? Are these distinct processes computed in the same or different brain areas? And if they are computed in the same region, how might the information be encoded in the neural population to permit the readout of these types of social information?

Ever since the early studies of patient HM, it has been clear that the hippocampus plays an important role in social memory^2^. Studies in mice have found that the hippocampus^3^, and in particular the dorsal CA2 region^4–7^ and the ventral CA1 region^8^, are crucial for social memory storage, consolidation, and recall through a dorsal CA2 to vCA1 circuit^7^. Neurons in dCA2 have been identified that change their firing to novel conspecifics^9,10^ and can distinguish the identity of two novel individuals^11^. Neurons in vCA1 have been found to increase their firing to social stimuli^12^ and selectively fire around familiar individuals^8^. While it is clear that these hippocampal regions encode social stimuli and are required for social memory, it remains unknown as to whether and how hippocampal activity can both support a generalized detection of familiarity while also providing the capacity to store a large number of detailed social episodic memories. Moreover, because hippocampal neurons also serve as place cells, encoding an animal’s position own position in its environment^13^, as well as the position of conspecifics it encounters^14,15^, it is uncertain as to how the hippocampus represents and disambiguates social and spatial variables to meet the conflicting demands of social memory.

Here we have addressed how the population activity of dorsal CA2 neurons may enable a generalized readout of familiarity while permitting the encoding of social/spatial episodic memories using a combination of calcium imaging and computational approaches. Based on our analysis of the activity of large populations of CA2 neurons as mice explored different combinations of novel and familiar individuals, we find that CA2 accommodates the competing demands of social memory by representing novel and familiar individuals in distinct geometric arrangements (i.e., the relationship between the neural population responses in distinct social situations in neural activity space). CA2 encodes novel animals in low-dimensional representations, which enable the identity of novel animals to be readily disentangled from their position. However, such representations suffer from a low memory capacity and so are not suitable for storing the vast information associated with past encounters with familiar animals. CA2 solves this problem by encoding familiar individuals in higher dimensional representations, increasing memory storage capacity at a modest cost to generalization across contexts. In addition, CA2 encodes familiarity separately from identity in an abstract, low-dimensional format. The degree of transformation in the dimensionality of novel and familiar representations in a given subject predicts the animal’s behavior in a separate social memory test requiring that animal to differentiate between a novel and familiar mouse. Together these results provide the first evidence that experiencedependent transformations in the geometry of neural social representations can enable a single neural population to both discriminate social novelty from familiarity while supporting the recollection of experiences with familiar individuals.

## Results

### Experimental approach and theoretical considerations of the geometry of social representations and its implications for social memory

Our goal in these experiments was to characterize how social identity, social familiarity, and spatial location are represented in conjunction with the neural activity of CA2 pyramidal neurons to support the underlying requirements of social memory. We used microendoscopic imaging of Ca^2+^ activity in dorsal CA2 pyramidal neurons in freely moving mice while they interacted with novel and/or familiar conspecifics confined to wire cup cages in an open arena. We then used computational and theoretical approaches based on machine learning techniques to probe the social and spatial information that was contained in CA2 neural representations to determine whether this information allowed social identity and spatial location to be disentangled and to infer the nature of the geometric structure of these representations.

We imaged a total population of 439 CA2 pyramidal neurons from 6 subject mice while they interacted freely with two conspecifics placed under wire cages at the two ends of an oval arena for 5 minutes (Fig. 1a, b, Suppl. Vid. 1). We then exchanged the positions of the two stimulus animals and allowed the subject mouse to explore the conspecifics for an additional 5 minutes to enable us to distinguish spatial from social activity (Fig. 1c). On different experimental days the subject mice were exposed to either two novel mice (N1 and N2), two familiar co-housed littermates (L1 and L2), or one novel mouse (N) and one littermate (L). When both mice had similar degrees of novelty or familiarity, the subjects showed no preference for exploration of either individual (Fig. 1d, Suppl. Fig. 1). We recorded Ca^2+^ signals during periods when the subject mouse was actively exploring one of the two stimulus mice through manual scoring of periods in which the head of the subject mouse was within an interaction zone extending 5 cm from the perimeter of the cup and the subject was engaged in sniffing behavior (Fig. 1d, f). We deconvolved the Ca^2+^ signals for each neuron to extract individual spikes that were grouped into 100-ms-long bins during interaction periods, labeled according to the identity (e.g., N1, N2) and position (left, right) of the social encounters. We then used a linear classifier to decode social identity and position from the population vectors of deconvolved calcium activity. Our analysis had the dual purpose of (1) quantifying the extent to which the social and spatial variables are encoded in CA2 population activity, and (2) revealing the structure of the population activity by analyzing its geometrical arrangement in neural activity space (Fig. 2).

**Figure 1.**
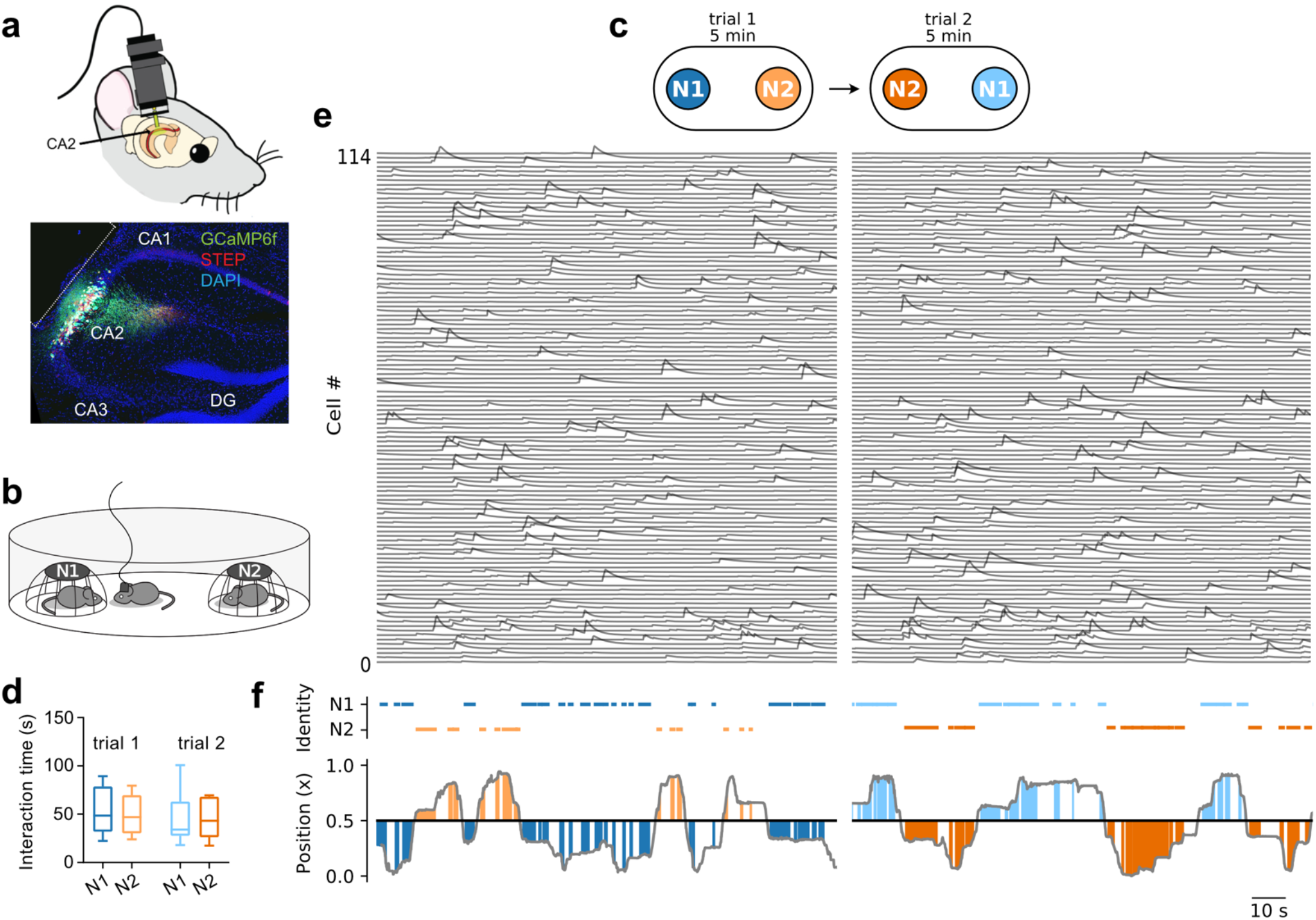
Experimental design. **a)** Six Amigo2-Cre subject mice were injected with a Cre-dependent GCaMP6f virus and implanted with a GRIN lens over dorsal CA2, enabling visualization of the activity of 439 CA2 pyramidal neurons. **b)** Implanted subjects ran through experiments in an oval arena with two stimulus mice (N1 and N2) under wire pencil cup cages, allowing interactions between the subject and stimulus mice. **c)** Subjects were first allowed to explore two novel individual stimulus mice in a five-minute trial. To distinguish social and spatial responses, the position of the stimulus mice were swapped in a second 5-minute trial. **d)** A plot of the total mean interaction time of the subjects with the stimulus mice in the two trials. No significant difference was observed for exploration of N1 or N2 in either trial: Two-way ANOVA for Partner x Trial F(1,5) = 0.0530, p=0.83. **e)** Example 114 simultaneously recorded deconvolved calcium traces across the two trials from a single subject mouse; **f)** Plot of the position of the subject along the axis defined by the center of the cups and identity of the interaction partner during the two trials. The colored lines on top and colored areas denote active interactions (sniffing) with either interaction partner. The four colors correspond to the four combinations of spatial (left versus right cup) and social (mouse N1 versus N2) variables.

We begin with a general consideration as to how the different possible geometries of neural representations under the conditions of our experiments impact the encoding and recall of social information. We then present theoretical results that illustrate the advantages and limitations of these different geometric arrangements. For illustration purposes, we consider the response of three CA2 neurons in the four conditions across the two trials of the experiment with two novel mice shown in Fig. 2a (trial 1: N1-left, N2-right; trial 2, N2-left, N1-right). In Figures 2b,c,d, each vector of activity of the three neurons in a given condition within a session is represented as a point. The precise location of each point will vary from vector to vector so that the neural responses to a given condition will form a cloud of points, and the responses in the four possible conditions of the experiment will form four clouds of points. The relative arrangement of these four clouds defines the *representational geometry* of the two variables (social identity and spatial position) that characterize the social episode.

**Figure 2.**
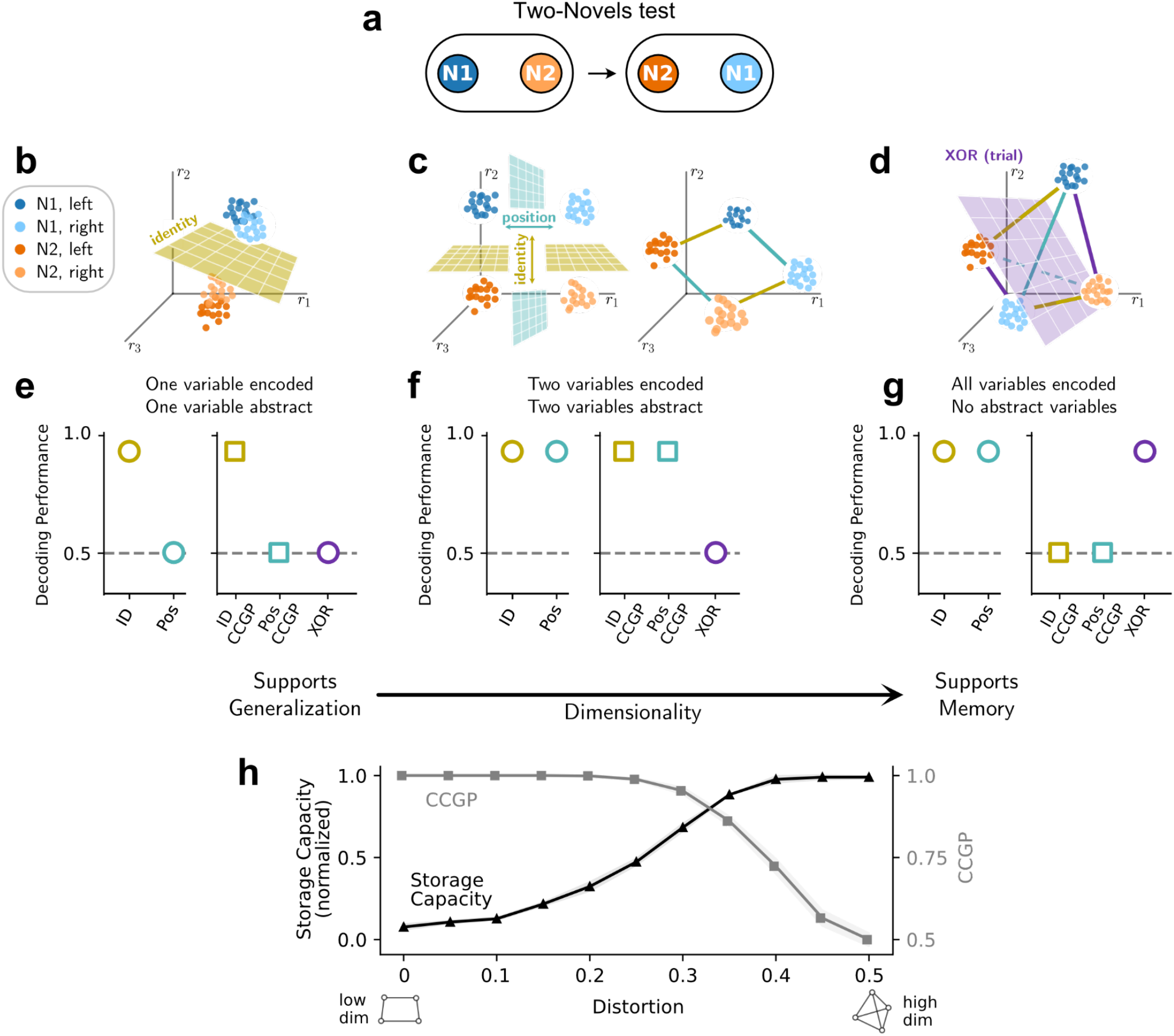
Neural recordings of dCA2 pyramidal neurons can elucidate the geometrical relationships between represented variables. **a**) Scheme of the two-novels experiment. **b-d)** Examples of geometrical arrangements of the coding of two variables (identity and position) with different dimensionality. Each point plots the firing rates of three neurons (*r_1_, r_2_, r_3_*) measured for a given color-coded condition. Because of variability in activity, multiple instances of a given condition results in a cloud of points. **b**) An example one-dimensional arrangement of the four conditions (N1-left, N2-left, N1-right, N2-right) in which identity but not position can be decoded. Identity is abstract to position in this orientation. A classifier plane (yellow) can readily separate the clouds of points to decode identity. **c**) Example twodimensional arrangements in which position and identity are encoded and disentangled. **Left,** the neurons are specialized, responding to either identity or position, resulting in a rectangular planar geometry. Two classifier planes (yellow and cyan) can readily separate the point clouds to decode identity and position. In addition, the classifiers generalize in that a classifier plane that optimally decodes identity when the animals are in the left cup will also decode identity when the animals are in the right cup (yellow planes), and vice versa. Similarly, the classifier trained to decode position when a given stimulus animal (e.g., N1) is in the two cups will also decode position when the other stimulus animal (e.g., N2) is in the two cups (cyan planes). **Right**, in this example, the neurons have mixed selectivity; they respond to both identity and position. However, they respond in a linear manner to the two variables, leading to a rotation of the points that maintains the rectangular-like planar 2-D geometry. This geometry also enables a linear classifier to perform with a high decoding accuracy and high generalization. Note the two geometries in **c** do not enable the decoding of points corresponding to the conditions represented at opposite corners of the rectangle-like shapes (N1-L & N2-R, N2-L & N1-R). This is termed the XOR dichotomy and corresponds to the conditions in the two trials. **d**) An example threedimensional, tetrahedral geometric arrangement in which position, identity, and XOR can be decoded by a linear classifier. Here, position and identity are entangled. **e-g)** different geometries in **b-d** have corresponding distinct fingerprints for their ability to decode identity, position and XOR, as well as for generalized decoding of identity and position as measured by CCGP. **h**) Tradeoff between storage capacity and CCGP in a simulation of a Hopfield recurrent neural network. A network of N neurons was trained to store and retrieve a set of patterns with geometrical dimensionality varying from L << N to N (see Methods and Supplementary Information). In this simulation, we used L=10, N=400. To vary the dimensionality of the patterns, we added a random distortion by flipping the value of each neuron in each pattern with a given probability ranging from 0 (L-dimensional, correlated activity) to 0.5 (N-dimensional, random uncorrelated patterns) with a step size of 0.05. Curves and points represent the average over n=10 model simulations. Storage capacity was normalized to range between 0 and 1.

Many different geometric arrangements are compatible with the previous observations that CA2 neurons encode identity and position^8^. However, these arrangements have distinct computational properties that lead to differences in the ability to read out the encoded social/spatial information. We used a binary linear classifier to explore how these geometries will impact the ability to read out the information contained within the population. The binary linear classifier, as opposed to non-linear classifiers, has two important properties. First, as illustrated in Fig. 2, it allows us to determine in a straightforward manner the geometric structure of the neural representations. Second, it has the neurobiological advantage of being readily implemented by downstream neurons that linearly sum their synaptic inputs from a population of CA2 neurons to produce an all-or-none output that reports the binary dichotomy under consideration (eg, social identity: N1 versus N2; or position: left versus right).

We first consider the geometric arrangement in the case that CA2 neurons only encode a single variable, social identity. In this case, pairs of point clouds that correspond to the same individual at different locations are clustered together, defining a simple one-dimensional (1D) object. A linear classifier trained to decode identity from neural activity identifies a plane that optimally separates the clouds of points. In this case, the classifier will perform well (Fig. 2e), as the cloud of population vectors determined for one identity (eg, N1, blue clouds in Fig. 2b) is well separated from that determined for the other one (N2, orange clouds in Fig. 2b). Importantly, a classifier trained to report identity when the animals are in the left cup (dark blue versus dark orange clouds in Fig. 2b) will be able to generalize to accurately report identity when the animals are in the right cup (light blue versus light orange clouds in Fig. 2b). This is termed the cross-condition generalization performance (CCGP)^16^. The simple, one-dimensional geometry shown in Fig. 2b yields a high decoding performance and a high CCGP for identity (Fig. 2e). Following Bernardi et al.^16^, we define this as an *abstract* representation of identity, as the neural responses to the different social identities do not depend on the position of the social interactions. However, such representations are of limited value for storing or encoding episodic memories as they do not encode any other variable than identity and, moreover, are incompatible with the observations that CA2 encodes both identity and position^8^.

Examples of more realistic neuronal populations that encode both identity and position are shown in Fig. 2c. In the case shown at the left in Fig. 2c, identity and position variables are encoded by two types of specialist neurons, each responsive to only one of the two variables. The clouds of activity vectors are arranged in a planar, rectangular-like shape (2-D), along the two axes corresponding to the firing rates of the two classes of neurons. In this case, separate linear classifiers can now decode both social identity and position (Fig. 2f). Moreover, both variables are abstract (high CCGP; Fig. 2f, Suppl. Vid. 2), since a classifier trained to decode identity in one pair of conditions (e.g., N1-right versus N2-right) can also decode identity in the other pair of conditions (e.g., N1-left versus N2-left), and a classifier trained to decode right from left position when both cups contain one animal (e.g., N1) can also decode right from left when the cup contains the other animal (eg, N2). This type of representation is called *disentangled* and is known to be important for generalization and compositionality^16–18^.

These same computational properties (generalization of both variables) can be maintained with neurons with linear mixed selectivity, i.e., that respond linearly to both identity and position. The activity of such neurons still results in a planar, rectangular-like geometric arrangement, but one that is rotated so that its edges are no longer aligned with the neural axes (Fig. 2c, right). However, the coding direction for a given variable is still parallel to the coding direction for that same variable across conditions so that a classifier trained to report identity at one location would generalize to the other location, and vice versa, hence yielding a high decoding accuracy and CCGP for both identity and position (Fig. 2f). Low-dimensional mixed-selectivity representations (Fig. 2c, right) have been observed in multiple brain areas^16,19^, and, as noted, they enable abstract coding of relevant variables.

A drawback of such low-dimensional representations is that they limit the number of variables that can be decoded. The four possible social/spatial combinations of our experiments can be arranged into three possible binary groupings, or dichotomies, that can be probed by a linear decoder. In addition to grouping the four combinations by social identity and position, a third grouping is possible in which neither of the social or spatial variables is consistent within a group (N1-left and N2-right versus N2-left and N1-right). This is termed the XOR dichotomy and corresponds to grouping the conditions according to trial. Neither of the 2D representations allows a linear classifier to separate these two pairs of point clouds as they lie at opposite corners of the rectangular representations.

In contrast to these relatively constrained low-dimensional (2D) geometries, the responses illustrated in Figure 2d show the least constrained, highest-dimensional (3D) representation possible with three neurons encoding four conditions. In this unconstrained case, the neural responses form a 3D tetrahedron. In this geometry, all three possible pairs of sub-groupings of the four conditions, including the XOR dichotomy, are linearly separable and individually decodable (Fig. 2g, Suppl. Vid. 3). This ability defines a representation with a high shattering dimensionality^16^. However, high-dimensional representations have a reduced capacity for generalization as the coding directions of each variable are no longer parallel across different conditions (Suppl. Vid. 4). As a result, the classifier plane that best separates a given pair of variables in one set of conditions (eg, N1-left versus N2-left) will not necessarily be able to separate the same variable in the other set of conditions (eg, N1-right versus N2-reight), resulting in a low CCGP (Fig. 2g). In summary, whereas low dimensional geometries provide for generalized decoding of identity/position and high CCGP without decoding of XOR, high dimensional geometries allow the decoding of a greater number of variables (including XOR) at the cost of generalization.

Of importance in considering the role of CA2 in social memory, dimensionality is directly related to the number of memories that can be stored in a recurrent neural network. For classical hippocampal-dependent episodic memory, the memory capacity is related to the number of distinct episodes that can be stored. As shown in Fig. 2h, and further developed with theoretical computations in the Supplementary Material, a low-dimensional representational geometry severely limits the memory storage capacity, where each memory is defined as a specific combination of variables (e.g., the encounter of an individual at a certain location). These limitations occur because low-dimensional representations have a greater correlation between the activity of different neurons compared to a high-dimensional representation, effectively reducing the number of independent neurons available to participate in memory storage. Conversely, a high dimensional geometry provides for a high memory capacity at the price of a reduced generalization capacity (Fig. 2e). Therefore, different geometries could satisfy different demands for social memory, with dimensionality controlling the tradeoff between generalization and memory storage capacity. In the next sections, we apply a linear classifier to characterize the geometry of social/spatial representations to determine whether and how familiarity alters the tradeoff between generalization and memory capacity to support different memory requirements in the hippocampus.

### Novel individuals are encoded in low-dimensional representations

We first examined the nature of CA2 social and spatial representations during the exploration of two novel mice by analyzing the activity in the entire pseudo-population of the 439 CA2 pyramidal neurons we recorded from all six mice run on the task. We first asked whether CA2 activity contained sufficient information to allow a linear classifier to decode the identity of the two mice by grouping in one class the activity recorded around mouse N1 in the two trials (N1-left in trial 1 and N1-right in trial 2) and in a second class the activity recorded around mouse N2 in the two trials (N2-right in trial 1 and N2-left in trial 2) as shown in Fig. 3a. Because we balanced the data so that the subject spent equal time around the left and right cups, decoding performance must reflect information about mouse identity without a confound of position. A linear classifier trained on a subset of data from both trials successfully decoded mouse identity using withheld data with a high accuracy that was significantly greater than chance (Fig. 3c).

**Figure 3.**
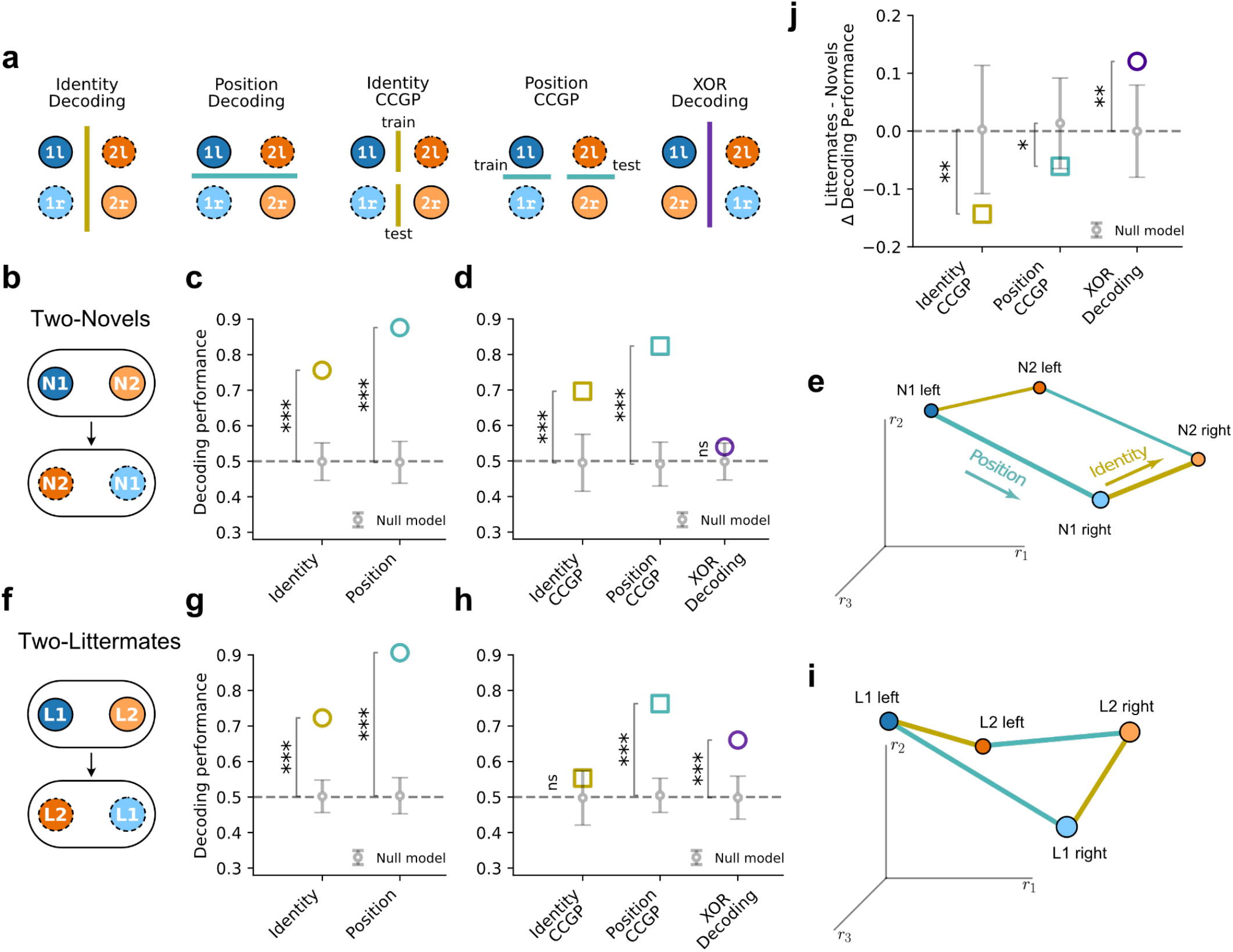
Novel and familiar identities are coded in different geometrical arrangements that support different memory requirements. **a**) Schemes for decoding identity, position, XOR (training and testing on data from all four conditions) and CCGP for identity and position (training on one pair of conditions and testing on the other). The numbers and colors indicate the identity of the stimulus animal under the cup (1, blue or 2, orange), the shade of the color indicates whether an animal is in the left cup (darker shades) or right cup (lighter shades) and the outline whether the data is from trial 1 (solid outline) or trial 2 (dashed outline). **b-e**) The scheme (**b**), decoding results (**c, d**) and proposed geometry (**e**) for the experiment with two novel stimulus mice. **c**) Open symbols show mean decoding performances compared to results of null model based on shuffled data (solid points and error bars, respectively). Novel mouse identity and position are decoded significantly above chance (id decoding = 0.76, null model = 0.50 ± 0.06; pos decoding = 0.88, null model = 0.50 ± 0.06). **d)** Identity CCGP and position CCGP are also significantly higher than the null model (id CCGP = 0.70, null model = 0.49 ± 0.04; pos CCGP = 0.82, null model = 0.49 ± 0.04). XOR coding does not differ from the null model (XOR decoding = 0.54, null model = 0.50 ± 0.06). **e**) Proposed low-dimensional geometry for social/spatial representations of two novel mice. **f-i**) The scheme (**f**), decoding results (**g, h**) and proposed geometry (**i**) for the experiment with two littermates as stimulus mice. **g**) Familiar mouse identity and position are decoded significantly better than chance (id decoding = 0.72, null model = 0.50 ± 0.06; pos decoding = 0.91, null model = 0.50 ± 0.06). **h)** Identity CCGP is not significantly greater than chance (id CCGP = 0.55, null model = 0.50 ± 0.05) while spatial CCGP and XOR decoding are significantly greater than chance (pos CCGP = 0.76, null model = 0.50 ± 0.03; XOR decoding = 0.66, null model = 0.50 ± 0.06). **i**) Proposed geometry for social/spatial representations of two familiar littermates. **j)** Difference in indicated decoding performance (Δ) in tests with two familiar stimulus mice compared to two novel stimulus mice. CCGP for identity and position were significantly greater with novel compared to familiar mice whereas XOR values were greater with familiar compared to novel mice (Δ identity CCGP = −0.14, null = 0.00 ± 0.06, p = 0.0042; Δ position CCGP = −0.06, null = 0.00 ± 0.04, p = 0.029; Δ XOR = 0.12, null = 0.00 ±0.04, p =0.0012). Values are reported as mean ± STD. Null model error bars show 2 STDs around the mean. P values are estimated from the z-score of the data values compared to the null model distributions. *p<0.05, **p<0.01, ***p<0.001.

Next, we asked whether CA2 activity contained sufficient information to decode whether a subject mouse was exploring the left versus right cup. In this case, we grouped activity data recorded from the two trials around the left cup in one class and around the right cup in a second class (N1-left and N2-left versus N2-right and N1-right), balancing data so that the mouse spent equal time exploring mouse N1 and N2 in the two classes. This removed social identity as a potential confound. The position classifier successfully decoded the left-right position of the subject mouse with a high accuracy that was significantly greater than chance (Fig. 3c). These two sets of results were not unexpected, given previous findings that a fraction of CA2 neurons can respond selectively around one novel mouse compared to another^8^ and that CA2 neurons also can serve as place cells, responding in specific locations even in the absence of another mouse^9–11,20,21^.

To determine whether social and spatial information was encoded by a specialized subpopulation of CA2 neurons (Fig. 2c, left, Suppl. Fig. 2a) or whether CA2 neurons had mixed selectivity (Fig. 2c, right, Suppl. Fig. 2b,c), we examined the weights assigned to each neuron by the linear classifiers trained to decode identity and position. We observed very few specialized neurons, no more than expected by random chance, as most neurons contributed to the decoding of both identity and position (Suppl. Fig. 2d-g). Moreover, the relative contributions of individual neurons to decoding identity and position were correlated, suggesting that neurons co-varied in the reliability of their social and spatial information (Suppl. Fig. 2d-g). From these results, we conclude that dCA2 neurons encode both spatial and social variables with mixed selectivity^22^.

To determine whether location and identity are represented in CA2 in a way that allows these variables to be readily disentangled and, hence, whether the variables are encoded in a generalizable or abstract format, we performed a CCGP analysis as discussed in Fig. 2. We first asked whether social identity was decodable independent of position by training a classifier to decode social identity when N1 and N2 were located in a given cup (e.g., the left cup) and testing whether that same classifier could decode identity when the mice were in the other cup (e.g., the right cup), as shown in Fig. 3a. Indeed, we found that CA2 neural activity supported a generalized decoding performance that was significantly greater than that expected by chance (Fig. 3d). Similarly, we also found significant generalized decoding of position, independent of the identity of the mice under the cup (Fig. 3d). These results suggest that identity and position of novel encounters are represented as abstract variables in CA2.

To further analyze the geometry of novel individuals, we probed the ability of a linear classifier to decode the XOR dichotomy, which corresponds to decoding trial number in our experiment (N1-left and N2-right from trial 1 versus N2-left and N1-right from trial 2). In contrast to the ability of CA2 representations to decode with high accuracy mouse identity and cup position, the classifier failed to decode the XOR condition above chance levels (Fig. 3d). Together, these results suggest that social encounters with two novel animals are represented in a low-dimensional, planar geometry (Fig. 3e).

### Familiar individuals are encoded in high-dimensional social-spatial representations

In contrast to meeting a novel individual for the first time, meeting a littermate is expected to both evoke a sense of familiarity and trigger the recall of specific episodes of prior social encounters, which are typically multi-dimensional experiences. To determine how familiarity may transform the social-spatial representational geometry in dCA2, we performed the above experimental protocol but now presenting two familiar littermates to the same set of subjects (Fig. 3f). Again, subjects explored the two littermates freely, showing no preference for either individual and no overall difference in exploration time when compared to the exploration times with two novel individuals (Suppl. Fig. 1). After 5 minutes of exploration, the positions of the cages were swapped for another 5-minute session to ensure balanced exploration times for position and identity variables (Fig. 3f). To allow for a direct comparison with data from the experiment with two novel mice, we further balanced the data so that each subject engaged in the same length of interaction for each condition across the two experiments. In addition, we used the same total number of neurons for determining the decoding performance across the two tests.

We found that CA2 population activity was able to decode identity and position with high accuracy during encounters with familiar individuals, at a level similar to that seen during encounters with novel individuals (Fig. 3g). Similar to the encounters with novel mice, CA2 neurons exhibited mixed selectivity for identity and position information during exploration of familiar littermates and we did not observe a significant fraction of neurons tuned to one or the other variable (Suppl. Fig. 2). However, compared to novel animals, we observed striking differences in the ability of CA2 representations of familiar individuals to generalize (disentangle) identity from position and to decode the XOR (trial) condition. First, we found that CCGP for familiar individual identity was not greater than chance levels (Fig. 3h). Second, whereas CCGP for position was still greater than chance, it was significantly less than the CCGP found during the exploration of the novel mice (Fig. 3h).

In contrast to the decrease in CCGP during the exploration of familiar animals, decoding performance for the XOR dichotomy was now significantly greater than chance levels (Fig. 3h), in contrast to what was observed with novel animals. As presented in Figure 2, a significant XOR decoding and a low CCGP is a signature of a higher-dimensional geometry (Fig. 3i). A direct comparison of the geometrical quantities observed during exploration of novel versus familiar mice showed that the observed differences were significantly greater than chance levels (Fig. 3j) estimated by a conservative null model for decoding difference (see Methods). These coordinated changes in geometrical quantities suggest that familiarity transforms the coding geometry of social encounters, increasing their representational dimensionality. An increase in dimensionality offers greater memory storage capacity, with a cost to generalization across contexts.

### Familiarity is represented as an abstract variable that generalizes across identity and spatial position

Our results so far focused on the representations of position and identity during social encounters and how the coding properties of these variables were modulated by familiarity. Although previous experiments using in vivo electrophysiological recordings showed that CA2 activity can be used to decode interactions with a novel versus a familiar animal^10^, those experiments did not determine whether decoding performance simply reflected the different social identities of the two individuals or whether CA2 encoded a generalized or abstract representation of novelty versus familiarity, independent of the particular identity of the individuals.

No prior study has addressed whether familiarity, identity, and spatial location are represented in CA2 in a way that allows these variables to be readily disentangled. Thus, we next investigated whether CA2 provides a general or abstract representation of social familiarity by allowing a subject to explore the arena with a novel conspecific and familiar littermate present under the two cups (Fig. 4a). Similar to the previous experiments, subjects were free to explore the two conspecifics for a first trial of 5 minutes. However, for the second trial the two stimulus mice were replaced by a new novel-littermate pair, with the positions of novel and familiar animals reversed (Fig. 4a). By analyzing the activity of CA2 pyramidal cells (438 neurons from 5 subjects) during active interactions with the two sets of novel and familiar individuals, we were able to test whether the coding of familiarity generalizes across spatial positions and social identities (Fig. 4b). We first confirmed that familiarity and position could be decoded from CA2 neural activity (Suppl. Fig. 3). We then trained a classifier to discriminate between a pair of novel and familiar animals located in the same cup across the two trials (eg, L1-left versus N2-left; see Fig. 4a, b) and tested whether this same classifier was capable of discriminating the different novel and familiar mice in the right cup (N1-right versus L2-right). We reasoned that if CA2 representations provided for an abstract coding of familiarity, we should observe a significant CCGP despite the difference in the specific identity of the novel and familiar animals and their different spatial locations.

**Figure 4.**
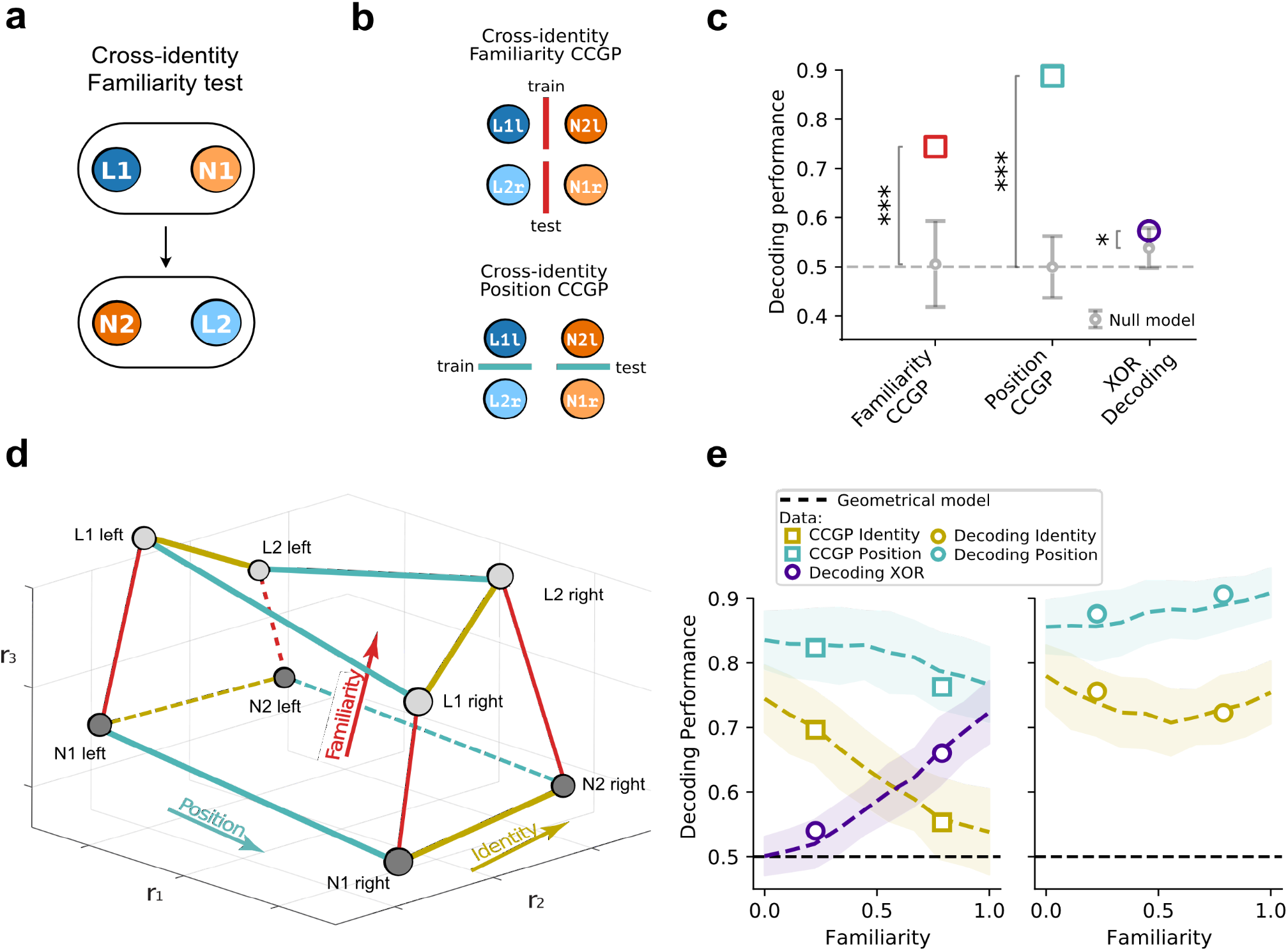
Degree of familiarity is encoded in a low-dimensional format disentangled from identity and position. **a**) Subject mice (n= 5 mice, 438 cells) were exposed to one novel stimulus mouse and one familiar stimulus mouse for a five-minute trial. The subject mice were then exposed to a second set of novel and familiar individuals in a second five-minute trial in which the position of the novel and familiar individual were swapped. **b**) Decoding scheme for familiarity CCGP and position CCGP. **c**) Familiarity and position CCGP and XOR decoding were significantly greater than the null model (familiarity CCGP = 0.74; null model = 0.51 ± 0.04; position CCGP = 0.89, null model = 0.50 ± 0.03; XOR decoding = 0.57, null model = 0.54 ± 0.02). **d**) Graphical representation of a geometrical model for encoding social and spatial information of novel and familiar mice by three example neurons (firing rates r1, r2, r3). Dark and light gray circles represent firing rates during specific combinations of social and spatial variables during interactions with novel and familiar animals, respectively. Increasing familiarity both shifts and distorts in neural firing space the planar, low-dimensional social-spatial representations of novel animals. **e**) A model based on the geometry depicted in **d** reproduces experimental observations (data from Fig. 3 is overlaid on predictions from the model. A best fit of the 6 parameters of the model reproduces our 10 decoding experimental data points, see Methods). Lines and shaded areas show mean ± SD calculated using 100 model simulations. Values are reported as mean ± STD. P values are estimated from the z-score of the data values compared to the null model distributions. Null model error bars show 2 STDs around the mean. * p<0.05, *** p<0.001.

Remarkably, despite the many variables that changed between the training and the testing conditions (trial, position, mouse identities), we found a high level of CCGP for discrimination of familiar versus novel individuals (Fig. 4c). Thus, CA2 representations implement an abstract code for familiarity that generalizes across social identities and spatial locations. Similarly, we also observed a high CCGP for position when we trained the classifier to discriminate left from right cups when they contained the two littermates and tested the ability of that classifier to discriminate left from right cups when they contained the novel mice. Consistently, the change in the activity of individual neurons from the novel to the familiar mouse was conserved across the two pairs of novel-familiar individuals (Suppl. Fig. 4c), suggesting that familiarity shifts the neural representation of conspecifics in a common direction in the neural space. The observed abstract code for familiarity did not rely on a global change in cell activity (Suppl. Fig. 4a, b). This analysis is consistent with a geometry in which the directions from novel to familiar are approximately the same for all individuals and all positions, allowing any linear readout trained on a pair of familiar and novel individuals to generalize to a new pair.

This result, along with our previous findings, allowed us to reconstruct a full geometrical picture of how identity, position, and familiarity of social encounters are represented in CA2 population activity. We aimed to encompass our experimental findings that: (1) novel encounters are represented in a low-dimensional, abstract geometry; (2) familiarization increases the dimensionality of representations, sacrificing generalization to increase the capacity for memory storage and retrieval; (3) familiarity acts on this representation by causing a more-or-less coordinated parallel shift in the two representations in a specific direction in the neural space, enabling the abstract encoding of familiarity. A schematic of a proposed geometric model that is consistent with our results is shown in Fig. 4d. Based on point 1, novel individuals were represented in a planar, rectangular-like geometry. Based on point 3, familiarity caused a progressive parallel shift in the geometry of novel animal representations. And based on point 2, familiarity also produced a distortion or twisting of the planar arrangement into an increasingly three-dimensional trapezoidal structure.

To test whether this geometric model could provide a reasonably precise quantitative description of our results, we calculated decoding performance and CCGP for the same variables that we studied in the experiments in an in-silico experiment. We generated a random Gaussian distribution of point clouds around each of the experimental conditions and measured decoding performance and CCGP, adjusting the parameters that characterized the geometry of the centroids (6 parameters, see Methods) to fit to our proposed geometrical experimental findings (decoding and CCGP in the two experiments of Fig. 3, for a total of 10 values). This geometric model was indeed able to provide a good fit to our decoding and CCGP results (Fig. 4e), showing that this geometric interpretation is quantitatively consistent with our data. This model also generated predictions for changes in activity that might occur with more intermediate degrees of familiarity than explored in the present experiments.

### The increase in representational dimensionality correlates with social memory behavioral performance

From the theoretical results presented in Fig. 2, we know that high dimensional representations are associated with a higher memory storage capacity. We therefore wondered whether the degree of modulation in representational geometry by familiarity in a subject mouse would reflect the behavior of this subject mouse when tasked with discriminating a novel from a familiar conspecific. If the geometric transformation reflects a memory process, we would expect that subjects with a smaller increase in representational dimensionality would show a lower behavioral performance, measured by the difference in time spent exploring a novel compared to a familiar individual.

To test this idea, we ran a social recognition test where the six subjects from the two-novel and two-littermate test were allowed to explore a novel and a familiar individual at the two ends of the oval arena (Fig. 5a), similar to the familiarity experiment described above (Fig. 4a). However, in the present case we presented the same pair of novel and familiar mice in the second trial. We confirmed that in a larger cohort (n=12 mice), subjects showed a preference for the novel over the familiar stimulus mouse during the initial presentation of the conspecifics in this task (Fig. 5b), consistent with prior reports^4,7^. We saw little behavioral preference for the novel compared to the familiar mouse in the second trial, presumably due to the decrease in novelty during the second exposure. We measured a combined behavioral discrimination index for both trials, defined as the time a subject spent exploring the novel mouse minus the time spent exploring the familiar mouse divided by the total social exploration time. We confirmed the importance of CA2 for this assay of social memory by using pharmacogenetic silencing of dCA2 to eliminate the behavioral preference for novelty in this task (Suppl. Fig. 5).

**Figure 5.**
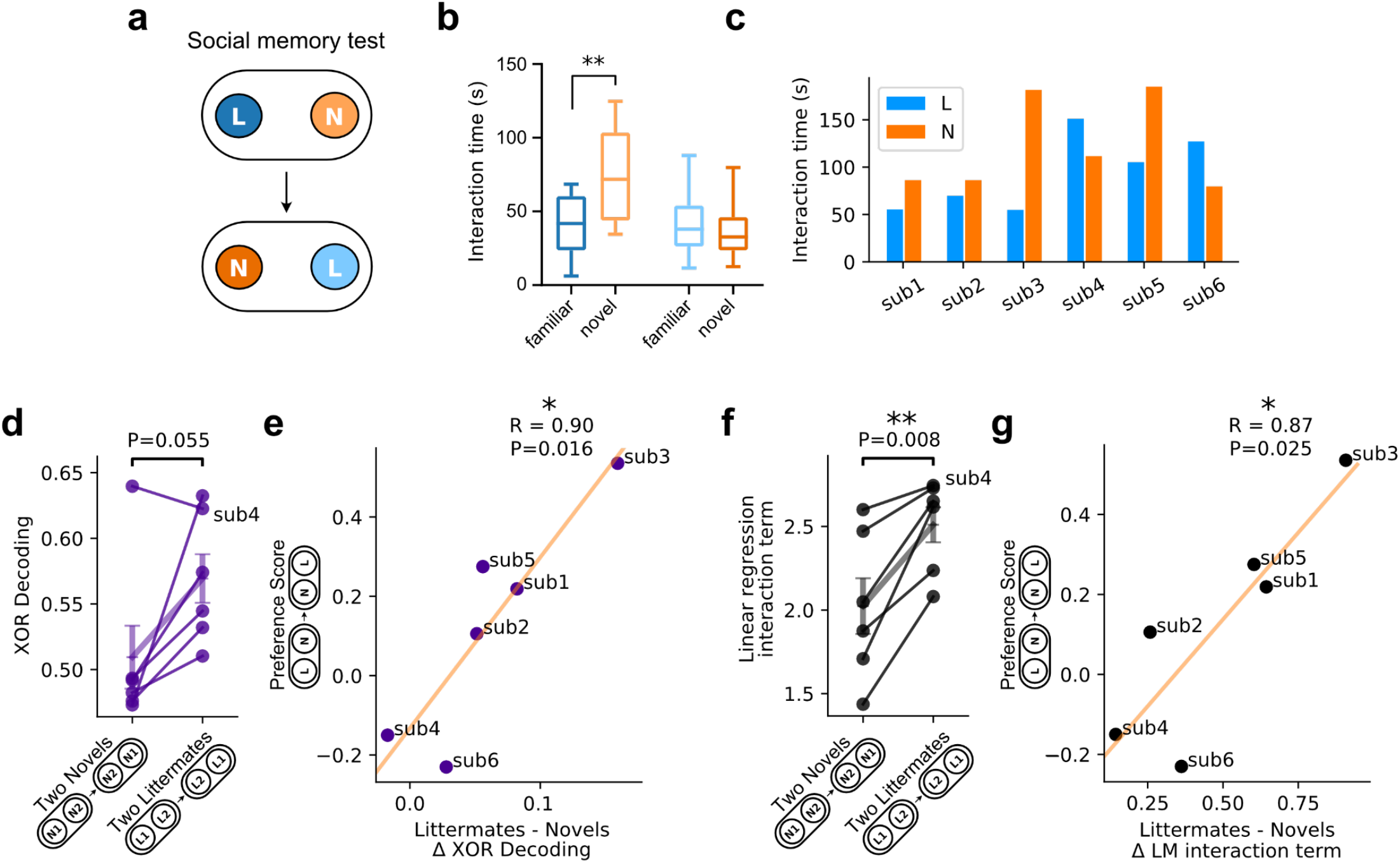
Magnitude of geometrical distortion in familiar and novel social/spatial representations is correlated with behavior in a separate social memory task. **a)** Subject mice performed a social recognition memory test in which they explored an arena with cups containing one novel and one familiar stimulus mouse across two five-minute trials (with positions swapped in the second trial). **b)** To validate that this test assesses social recognition, a total of 12 mice underwent the test, including the 6 implanted subjects from Figure 3. The subjects spent on average significantly greater time exploring the novel compared to the familiar stimulus mouse in trial 1 but not in trial 2 (two-way ANOVA: Interaction Partner x Trial, F(1,11) = 9.208, p=0.011. Šídák’s multiple comparisons test: trial 1 p=0.0085; trial 2 p=0.75). **c)** Total interaction time with the novel and with the littermate across the two trials, from the subset of 6 mice that were run in two-novels and two-littermates experiments. **d)** XOR decoding during trials with two novel and two familiar stimulus mice for individual subject mice. Mean XOR decoding value was significantly different from chance level of 0.5 with familiar mice (0.57 ± 0.045). There was a strong trend for a greater mean XOR with familiar compared to novel stimulus mice that did not reach significance (two-novel mice: XOR mean = 0.51 ± 0.03 SEM; two familiar mice: XOR mean = 0.57 ± 0.02 SEM; n=6, Student’s paired t-test, p=0.055). Points show values for the individual subjects. Vertical and horizontal lines show mean ± SEM. **e)** The magnitude of the difference in XOR decoding performance for individual animals was significantly correlated with behavioral preference for the novel compared to the familiar mouse in the separate social memory test of panel **a** (r=0.90, p=0.016). **f)** Interaction term from ANOVA performed for CA2 firing rates as function of mouse identity and position with two novel or two familiar stimulus mice. Mean interaction term value (position x identity) was significantly greater with two familiar mice (2.51 ± 0.12 SEM) compared to two novel mice (2.02 ± 0.18 SEM; Student’s paired t-test, p=0.0084, n=6 mice). Points show values for the individual subjects. Vertical and horizontal lines show mean ± SEM. **g)** The interaction term was correlated strongly with behavior preference for the novel over the familiar individual in the social memory test of panel a (r=0.87, p=0.025). *p<0.05, **p<0.01.

Because the XOR is a sensitive measure of the transition from low to high dimensional geometries^16^, we compared the change in XOR decoding observed during interactions with two littermates versus two novel mice (Fig. 3) with the behavioral discrimination index measured during exploration of the novel mouse and littermate in the experiment of Fig. 5a for each of the six subjects. Strikingly, we observed a very strong positive correlation between these two measures, with the magnitude of the change in XOR during interactions with two familiar compared to two novel individuals (R=0.90, p=0.016, Fig. 5e). This correlation is particularly noteworthy as the XOR and behavioral results were obtained using distinct sets of novel and familiar mice encountered 2-3 weeks apart. In contrast, neither the decoding performance for identity nor position had a significant relationship with behavior (Suppl. Fig. 6).

As individual neurons of high-dimensional representations respond in a non-linear manner to changes in more than one variable, we obtained a second measure of dimensionality by using ANOVA to fit a linear model of the response of individual neurons to the two stimulus variables (position and identity). The reported mean non-linear interaction term between the two variables (position x identity) across neurons within individual subjects thus provides a measure of dimensionality^17^. Individual subjects showed a consistently greater interaction term for representations observed during exploration of two familiar individuals compared to values during exploration of two familiar individuals (Fig. 4f). Importantly, the magnitude of the increase in the mean interaction term was also strongly correlated with behavioral performance in the social recognition test (R=0.87, p=0.025, Fig. 5g), further strengthening the link between the strength of social memory and representational dimensionality.

## Discussion

Our experiments and analyses, based on large-scale calcium imaging of hippocampal dCA2 pyramidal neuron activity from mice engaged in social encounters, demonstrate that dCA2 simultaneously meets the demands of social recognition memory by representing novel and familiar animals in distinct geometries optimized for generalization or higher memory capacity. We focused on dCA2 because of its prominent role in the encoding, consolidation, and recall of social memory^4,5,7^. Previous studies using in vivo recordings found that dCA2 neurons respond during social interactions^10,11,23^ and can distinguish between a novel and a familiar animal^10^. At the same time, CA2 neurons have been found to act as place cells, firing in particular regions of an environment either in the absence or presence of a conspecific, although CA2 place fields are less precise and stable than those of its CA1 and CA3 neighbors^9,10,21^. Despite these insights, to date, it has been unclear how dCA2 representations of social/spatial information support the multiple requirements of social memory, including the ready discrimination of social novelty versus familiarity, along with the storage and recall of social episodic memories formed during encounters that lead to familiarization with a given conspecific.

Although our results indicate that dCA2 neurons encode the individual identities of novel and familiar animals with similar degrees of precision, we found that the representations differ in two important respects. First, the social/spatial representations of novel individuals provided for a greater degree of generalized or abstract readout of social and spatial information assessed by the cross-condition generalized performance (CCGP). This further indicates that representations of social identity and spatial position were disentangled for novel but not familiar individuals. Second, social/spatial representations of familiar individuals, but not novel individuals, allowed for the decoding of the XOR dichotomy. Of importance, the social/spatial representations of novel and familiar individuals also support the abstract coding of familiarity, independent of the specific identities or positions of the familiar and novel individuals.

A simple geometrical model was able to capture the key elements of our findings. According to the model, novel individuals and their spatial location are represented in a low-dimensional, planar geometry, which provides for a generalized, or abstract, disentangled representation of social identity and position. The increased familiarity of a co-housed littermate, presumably resulting from multiple encounters with the same individual over an extended period of time, is associated with two distinct changes in the geometry of social representations. First, familiarity causes a rigid translation that produces a roughly parallel shift in the lowdimensional representations of novel animals in neural activity space. This allows familiarity to be encoded in an abstract format, similar to what has recently been observed in the human medial temporal lobe for image recognition^24^.

Second, the shift is accompanied by a distortion that increases the dimensionality of the social and spatial representations, allowing for an increased memory capacity. These two transformations are observed in multiple animals and are strongly correlated to each other. Indeed, we found that the extent of the change in dimensionality of familiar versus novel representations is strongly correlated to a subject’s behavioral performance in a social novelty memory test (Fig. 5). What is particularly striking is that the social novelty test and the neural recordings of social representations of pairs of novel and familiar individuals were performed on separate weeks with separate novel animals, suggesting the robustness of the difference in geometries on social recognition behavior. Our results thus indicate that the distinct geometric arrangements of familiar and novel animal representations provide a neural mechanism for how the brain may rapidly distinguish a novel from a familiar individual while enabling the storage of detailed information about social experiences with familiar animals. These transformations likely reflect a common process of long-term synaptic plasticity, although the precise mechanisms responsible for the changes remain to be determined. Through such a transformation, a single brain region is able to support the two key components of social recognition memory: social familiarity and social identity recollection.

Interestingly, our results show similar aspects to recent descriptions of representational geometry for familiar and novel faces in the monkey inferotemporal (IT) cortex^25^. Similar to our findings, the representations for novel faces are low dimensional (see ^19^). At short latencies, the dimensionality of familiar and unfamiliar face representations is similar, with the two geometries related by a simple translation. In contrast, at longer latencies, the geometry of familiar representations becomes distorted.

Our results provide the first demonstration, to our knowledge, that experience-dependent changes in representational geometry are associated with a behaviorally relevant differential cognitive processing of novel and familiar individuals. The balance of generalization and memory achieved with these different representations is likely an important feature that guides the encoding of complex social relationships to form a cognitive map of social space. Such coding may both be a product of and be required for navigating complex social behaviors, such as pair bonding, social aggression, and the creation of social dominance hierarchies, which will be of interest to explore in future studies. Moreover, as abnormalities in social cognition are a hallmark of various psychiatric disorders, it will be of further interest to determine whether deficits in social memory in various mouse models may be reflected in a loss of plasticity in the geometry of social/spatial representations.

The distortion that we observed for familiar encounters is likely to be an important universal component of efficient memory storage that goes beyond social memory. The idea that episodic memories should be “re-coded” to be stored more efficiently dates back to the studies of David Marr^26^. Re-coding has been the main idea behind the random non-linear transformations proposed in several studies^27–30^. These transformations were designed to generate well-separated representations of different memories, a function usually referred to as “pattern separation”. Recent models proposed that pattern separation is a signature of a process of memory compression, used by the hippocampus to generate more efficient decorrelated representations^31–33^. In all these cases, the transformations increase the dimensionality of the representations, similar to what we observed in CA2 activity during interactions with littermates. Therefore, the reported increase of dimensionality could be the signature of efficient memory encoding, an effect that we predict should be seen in many other situations that involve different types of memories.

## Author contributions

L.M.B. and S.A.S conceived the project and designed the experiments with input from L.P. and S.F., L.M.B. and S.I. collected the data, and L.M.B. and L.P. analyzed the data with guidance from S.F. The data was interpreted by L.M.B., L.P., S.A.S, and S.F., who wrote the paper with feedback from S.I.

## Data and materials availability

All data and materials are available from the lead corresponding author with reasonable request.

## Code availability

All custom code used for data collection and analysis are available from the lead corresponding author with reasonable request.

## Acknowledgments

We thank T. Tabachnik for help designing and obtaining the oval arena; R. Sahai, K. Lewis, and A. Fisher for assistance in obtaining immunofluorescent images and implementing DeepLabCut; D. Salzman, R. Hen, S. Hassan, P. Kassraian-Fard, and A. Villegas for critical discussions and comments on the manuscript. We are grateful to R. Nogueira, V. Fascianelli, S. Muscinelli, and J. Minxha for many valuable and knowledgeable discussions, and to MG. Posani for help with graphical illustrations.

## Methods

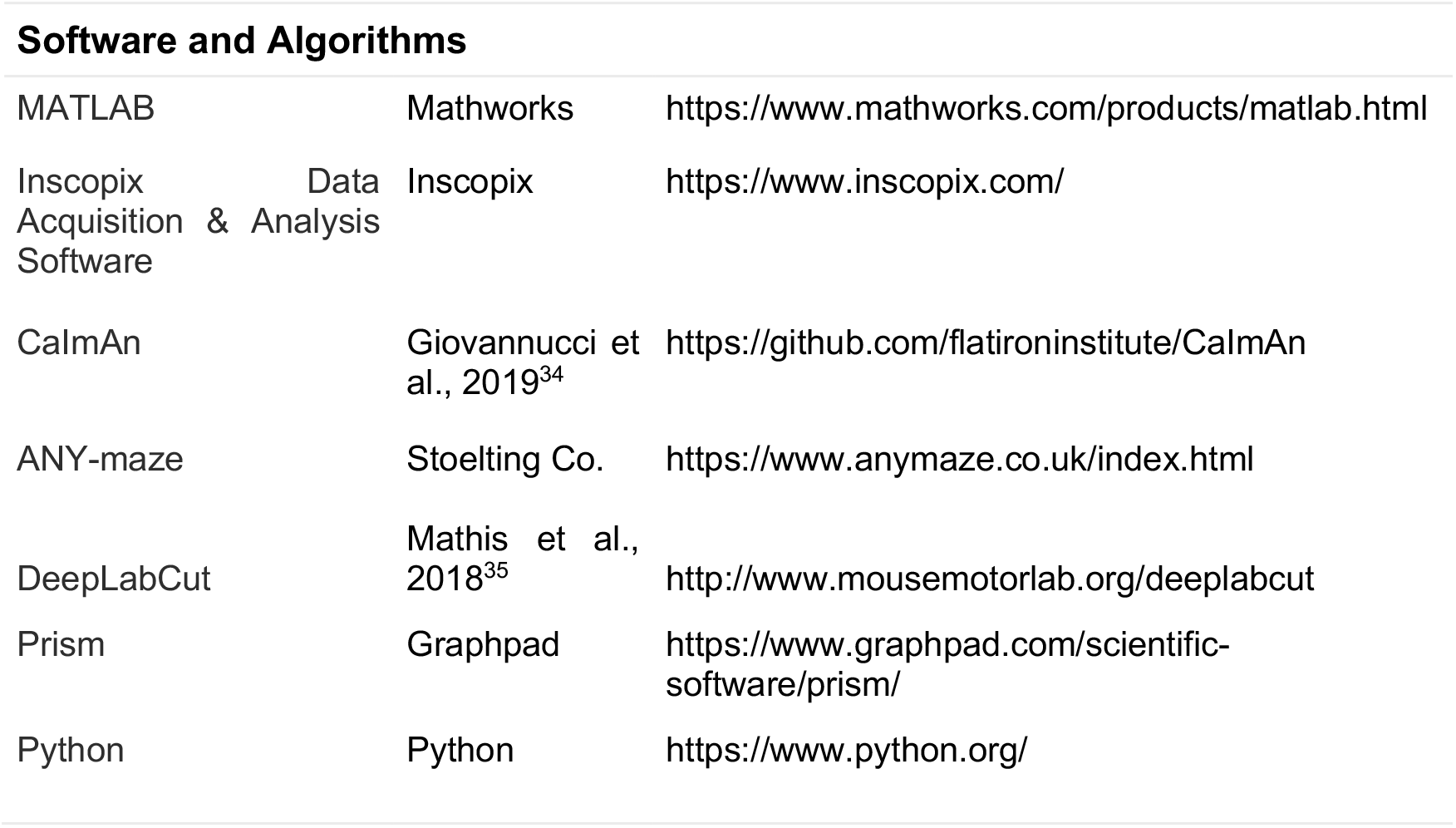

### Viral injection and GRIN lens implantation

#### Calcium imaging

A volume of 200 nL AAV2/1.syn.FLEX.GcaMP6f.WPRE.SV40 virus (titer: 6.5 x 10^11^ pp/mL, Penn Vector Core) was injected at a rate of 150 nL/min into the right hemisphere above dorsal hippocampal CA2 using stereotactic coordinates: AP −2.0 mm, ML +1.8 mm, DV −1.7 mm from bregma of 3-6 month-old male heterozygous Amigo2-Cre (Cre^+/-^) mice. Three weeks following injection, a 1.2 mm diameter circular craniotomy was centered at the following coordinates: AP −2.0 mm, ML +2.5 mm. We inserted a GRIN lens (Inscopix, 1.0 mm diameter, 4.0 mm length) into the craniotomy at a depth of −1.4 to −1.5 mm relative to bregma at a 10° angle from the midline, so that the lens was parallel to the CA2 cell body layer. The Inscopix Proview system imaged cells during implantation to adjust the position of the lens to optimize visible fluorescence. Kwik-sil was placed around the craniotomy and the lens secured in place using Metabond dental cement. The top of the Proview lens cuff was filled with Kwik-cast to protect the lens. Mice were housed with littermates for one week before a plastic baseplate was placed over the lens and secured with Metabond dental cement. The baseplate and microscope were placed over the lens and the position was adjusted until cells were maximally in focus.

#### Pharmacogenetic silencing of CA2

We injected 8 Amigo2-Cre^-/-^ (controls) and 12 Amigo2-Cre^+/-^ male mice in dCA2 with a Cre-dependent virus expressing the inhibitory hM4Di designer receptor exclusively activated by designer drugs (iDREADD), AAV2/8 hSyn.DIO.hM4D(Gi)-mCherry. 200 nL of virus (1.9×10^12^ pp/mL) was injected into dCA2 bilaterally using the following coordinates: anteroposterior (AP) −2.0mm, mediolateral (ML) +/-1.8mm, dorsoventral (DV) −1.7mm.

#### Immunofluorescent Labeling & Imaging

We perfused mice at the end of the experiments using saline followed by 4% PFA in ice-cold PBS. Brains were extracted and incubated in 4% PFA overnight. Brains were sliced in coronal orientation with thickness of 60 μm using a Leica VT1000S vibratome. Sections were permeabilized and blocked for 1 hour with 5% goat serum and 0.4% Triton-X in PBS at room temperature. Sections were incubated overnight with a CA2 marker primary antibody, either pcp4 (1:300, rabbit anti-pcp4 #HPA005792, Sigma-Aldrich) or STEP (1:1000, mouse anti-STEP # 4396, Cell Signaling Technology) at 4°C in 0.1% Triton-X in PBS plus 5% goat serum. The following day, slices were washed with PBS three times for 10 minutes in PBS and incubated with secondary antibodies (respectively: 1:500 goat anti-rabbit IgG, Life Technologies, or 1:500 goat anti-mouse IgG1, Life Technologies) for three hours. Slices were again washed three times in PBS for 10 minutes/wash. DAPI (ThermoFisher Scientific, #D1306) staining was applied at 1:1000 for 15 minutes in PBS at room temperature prior to mounting. Slices were mounted using Fluoromount (Sigma-Aldrich) and imaged using Zeiss LSM 700 confocal microscope.

### Extraction of Calcium Signals

#### Data Acquisition, Preprocessing and Motion-correction

On the day of the experiment, mice were moved to the behavior room and subject mice and littermates were separated into holding cages. Mice were allowed to acclimate to the environment for 30 minutes. An nVista 3.0 Inscopix miniaturized microscope was inserted into the baseplate and used to record calcium fluorescence from dCA2 pyramidal neurons during social and non-social behavior using Inscopix data acquisition software (20 frames per second, 50-ms exposure, 0.2-0.3 mW/mm^2^ EX-LED). The working distance between the microscope objective and the lens was adjusted to maximize cell focus, and this distance was maintained between trials and from session to session. To align behavior and calcium videos, a 5V TTL pulse from an Ami-2 Optogenetic interface triggered calcium recordings through Anymaze software at the start of each trial along with a behavior video recording. Behavior recordings were collected at a rate of 20 Hz. The raw videos from separate sessions were concatenated and then run through Inscopix Data Analysis software. Videos were preprocessed to correct defective pixels and 4x spatially down-sampled. Background fluorescence was removed using a spatial band-pass filter and fluorescence videos were motion-corrected using the Inscopix motion correction algorithm. The preprocessed and motion corrected tiff files were then exported for cell identification and signal deconvolution.

#### Segmentation and ROI Identification

Cell regions-of-interest (ROIs) were identified using the Python CaImAn package for large-scale calcium imaging data. The spatial footprints and deconvolved signal for the active sources (ROIs) were extracted using CNMFe^36^, and then the scaled raw traces and spatial footprints were exported to Matlab. We used a custom GUI to evaluate individual ROIs and spatial footprints, and those with non-spherical or non-oval shapes caused by motion artifacts were excluded from analysis. We detrended the raw traces over a window of 50 s using custom scripts. Finally, the computed traces, separated by session, were deconvolved using the OASIS algorithm for nonnegative signal deconvolution (baseline = trace median, noise = trace MAD, spike thresholds = 2x MAD).

### Behavior

#### Calcium recordings

We imaged dCA2 pyramidal neurons in a total of nine Amigo2-Cre heterozygous male mice in multiple tests probing social recognition and memory. Prior to the first test, mice were handled and habituated for three days on the following schedule: Handling (day 1), handling, exposure to oval arena for 15 minutes (day 2), handling, exposure to holding cage for 30 minutes, scruffing/insertion of the microscope, and to the oval arena for 15 minutes with microscope inserted (day 3). Mice were additionally habituated in the oval arena to empty cups for 10 minutes. No changes in subject mouse behavior, including during social interaction, were observed compared to wild-type controls.

In each test, subject mice were placed into an oval arena that consisted of two half-circles with radius 15 cm connected to a central square area with length of 30 cm (total dimensions: length 60 cm, width 30 cm, height 45 cm). Wire pencil cups (radius 5 cm) were placed 10 cm from the two ends of the arena along the midline and will hereafter be referred to as left cup and right cup. Stimulus mice were placed underneath the cups as described for each test. Between consecutive trials, subject mice were removed to a holding cage to which they had been previously habituated for approximately 2 minutes while the oval arena was cleaned with 70% alcohol wipes to remove any olfactory cues, wiped with paper towels, cleaned with water, and then wiped with paper towels until dry. The cups with or without stimulus mice were re-introduced to the arena, and finally the subject mouse was reintroduced into the arena and the trial initiated in ANY-maze. The position of the two stimulus mice were randomized to the left or right cups in the first trial, and the positions then swapped in the second trial. Stimulus mice were age- and sex-matched to subject mice (3-6 months old).

In each trial, the subject mouse was free to explore the arena. Periods of interaction with cups or conspecifics in the arena, defined as times when the subject’s head was oriented towards the center of the cup within a zone equal to 2x the cup radius (10 cm), while the subject was actively sniffing, were manually scored. In a minority of tests and trials, the subject mouse climbed on top of the wire pencil cups. In these cases, the period atop the cup was excluded from analysis. The behavior videos were run through a deep neural network trained using DeepLabCut to recognize the position of the mouse head and body, as well as location of the objects placed in the arena. Errors in the DeepLabCut output were corrected using an automated custom Matlab script.

#### Interaction with mice with similar degrees of novelty or familiarity

Six subject mice (neuron n=439) were exposed to two novel mice using three 5-min trials: habituation trial, two empty cups; trial 1, two novel mice; trial 2, the same two novel mice with positions swapped (Fig. 1c). The same six subject mice (neuron n=595) were exposed to two familiar littermates using three 5-min trials: habituation trial, two empty cups; trial 1, two familiar littermates in the cups; trial 2, the same familiar littermates with positions swapped (Fig. 3f). Subsequent tests were run at least one week apart.

#### Social Novelty Recognition Test

Five subject mice (neuron n=438) underwent the following three 5-min trials: habituation trial, two empty cups; trial 1, novel mouse 1 and familiar littermate 1; trial 2, novel mouse 2 and familiar littermate 2, with novel/familiar animal positions swapped relative to trial 1 (Fig. 4a).

#### Familiar versus novel mouse recognition test

Twelve subject mice underwent the following three 5-min trials: habituation trial, two empty cups (left and right); trial 1, novel mouse and familiar littermate in the two cups; trial 2, same novel mouse and familiar littermate with positions swapped (Fig. 5a). Of these twelve subject mice, six were additionally run in the two-novels and two-littermates tests as described above.

#### Behavior Statistical Analysis

To determine whether there were significant differences in the interaction times of the subject mouse with different social stimuli, we ran a two-way ANOVA of trial and interaction partner with repeated measures for both factors using Graphpad Prism software (version 9.0.1). Šidák’s multiple comparisons test was used post-hoc to determine significant differences between interaction partners. Statistical significance was defined as p < 0.05. As a measure of preference for one interaction partner (B) against the other (A), in Figure 5e and 5g, we calculated the social preference score defined as:

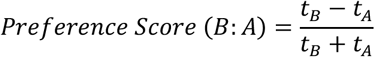

Where *t_A_* and *t_B_* are the length of time the subject mouse interacted with mouse A and mouse B across both social interaction trials.

#### Effect of CA2 silencing on social memory

Three weeks after iDREADD viral injection, Amigo2-Cre heterozygous mice (n=12) and wild-type littermates (n=8) were habituated to IP injection for four days. On the third and fourth day, mice were additionally habituated to the same oval arena used in calcium recording experiments for 5 minutes and to an individual holding cage for 30 minutes. On the fifth day, mice were moved to the experimental room and allowed to acclimate to the environment for 30 minutes in their individual holding cages. Mice were then injected intraperitoneally 30 minutes prior to testing with 10 mg/kg clozapine-n-oxide (CNO), the ligand for the iDREADD receptors, to reduce CA2 activity.

30-minutes post-injection, subject mice were run through two 5-minute learning trials in the oval arena: trial 1, novel mouse 1 and novel mouse 2 in the two cups; trial 2, the same two mice with positions swapped. In between each trial, the subject mouse was returned to the holding cage for approximately 2 minutes. Following trial 2, the subject mouse was returned to its holding cage. After a two-hour interval, the subject mouse was returned to the arena for a memory recall trial: trial 3, one of the previously encountered mice in the learning trials (e.g. novel 1, now familiar 1) and a third previously unencountered novel mouse (novel 3). The behavior videos were manually scored for interactions, defined by the same criteria as those applied during calcium imaging behavior, by an investigator blinded to the identities of the subject mice and the individuals under the cups. Memory recall was assessed by the greater interaction time with novel 3 compared to the previously encountered mouse, using the same statistical analysis described to determine social memory above.

### Population decoding analysis

The decoding analysis was performed using a linear classifier based on a support vector machine with custom-written Python scripts based on the scikit-learn SVC package^37^.

#### Data labeling

For each subject and session, we selected neural data corresponding to periods in which the subject was actively interacting with one of the two cups. We then divided the neural recordings into 100 ms time bins and labeled them according to whether the subject was interacting with the left or right cup and to the identity of the animal under the cup (labeled as *#1* or *#2*). In each test there were always two trials, with the positions of animals swapped in trials 1 and 2. Thus, for each test there were a total of 4 social/spatial conditions [mouse 1 on left (*#*1-left), mouse 1 on right (#1,-right), mouse 2 on left (*#*2-left), mouse 2 on right (*#*2-right)]. We then divided the four conditions into binary dichotomies (class 0 and class 1) according to the variable we wished to decode. For example, *social stimulus identity* was decoded by grouping firing data around the familiar animal as class 0 (#1-left & #1-right) and grouping firing activity around the novel animal as class 1 (#2-left & #2-right). We decoded *stimulus position* by grouping firing activity around the left cup as class 0 (#1-left & #2-left) and grouping firing activity around the right cup as class 1 (#1-right & #2-right). For XOR decoding, we grouped together conditions that have no identity or position values in common, defining two classes that incidentally correspond to trial 1 and trial 2 of our experimental setup: (#1-left & #2-right) as class 0 and (#1-right & #2-left) as class 1.

#### Cross-validation and pseudo-simultaneous population activity

For each subject and session, we divided data from each class of conditions (0 and 1) into training and test *pseudo-trials*, which each trial defined by a bout of interaction, with bout duration lasting from the beginning to end of a given interaction. Bout durations lasting longer than 1 s were split into multiple 1-s-long pseudo-trials. We randomly selected 75% of pseudo-trials for training a classifier and the remaining 25% were used for testing decoding performance. We next constructed a set of pseudo-population activity vectors from the training and testing datasets from a given animal by dividing each pseudo-trial into 100-ms bins, with each bin having its associated population activity vector containing the mean event rate observed during that time bin for each neuron recorded. We then randomly sampled *q* population vectors (where q=5 unless otherwise noted) from the training data set of each subject and concatenated them to form a single ņn-long vector, where *n* is the total number of recorded neurons in a given subject. This procedure was repeated *T* = 2*qn* times to create a training data set of pseudo-population firing rate vectors. We then followed the same procedure to build the pseudo-population testing data vectors, by sampling population vectors from the testing data set of each subject. In some cases, we performed decoding analysis on data from all N neurons from all animals tested in a given behavioral task. In this case, we randomly sampled *q* population vectors from the training data set for each individual animal. Next, we concatenated those extended population vectors into one pseudo-simultaneous qM-long vector. We repeated this process sampling successive sets of random population vectors for a total of *T* = 2*qN* pseudo-simultaneous training set vectors. We then repeated this process to obtain the testing data set vectors. To disentangle the selectivity to position and stimulus identity, which are correlated variables, the sampling procedure described above was performed in a balanced way so that each condition within each class was equally represented in the training and testing pseudo-simultaneous data set (e.g., for identity decoding: balancing #1-right and #1-left for class 0 and balancing #2-right and #2-left for class 1). The pseudo-simultaneous training data set was then used to train a SVM linear classifier, which was tested on the pseudo-simultaneous testing data set to assess the decoding performance as the fraction of correctly classified pseudo simultaneous vectors. The whole procedure, from training-testing division to performance assessment, was repeated for *k* = 20 times to implement a k-fold cross-validation scheme, taking the mean score (μ_*data*_) as the estimated performance value of the decoding procedure. To allow for a meaningful comparison of decoding results across experiments, only subjects that explored the four conditions (#1-left, #1-right, #2-left, #2-right) for a minimum of 3 s each, divided into a minimum of 4 pseudo-trials, in all three experiments (two-novels, two-familiars, novel-familiar) were used in the decoding analysis.

#### Null model and p-value

We tested the decoding performance obtained by the cross-validated procedure described above against a null model where the labels (*0* and *1* as defined above) of pseudo-trials were randomly shuffled. After each shuffling, the same cross-validation procedure was repeated, obtaining a null-model value for decoding performance. We repeated the shuffling *n_null_* times to obtain a distribution of null model performance values, yielding a mean null decoding performance (<μ_*null*_>) and standard deviation of the null distribution (σ_*null*_). The *p* value was then derived from the z-score of the performance computed on data compared to the distribution of *n_null_* null-model values: *z* = [μ_*data*_ − <μ_*null*_ >]/σ_*null*_.

#### Multi-selectivity analysis

We performed the following analysis to assess whether decoding of position and identity was primarily driven by cells specialized for one of the two variables (social identity or spatial location of the stimulus). For a given variable *X* and each cell *i* we identified the coding importance of each individual cell, defined as 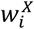, as the absolute value of its average decoding weight over *k* cross validation folds:

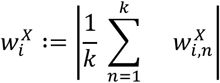

Where 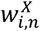 is the SVM decoding weight of cell *i* cell in the *n*-th cross validation fold. We then obtained a relative coding importance by normalizing the vector of all the values across the recorded population of cells:

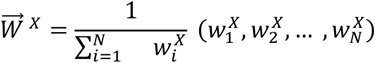

We denoted these vectors as 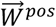, for position decoding, and 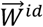 for social familiarity/identity decoding. If the recorded population is specialized, neurons that encode position will not encode identity, and vice-versa. Therefore, a population of specialized neurons will be characterized by anti-correlated values of 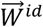 and 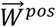 (Supp. Fig. 2 a). On the other hand, if neurons are not specialized (mixed selectivity), we expect no relationship between identity and position coding, resulting in a null correlation between 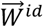 and 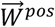 (Suppl. Fig. 2b). A third possibility is that neurons are not specialized, but information is unevenly distributed across the population. In this case, neurons will typically encode neither or both variables, resulting in a positive correlation between 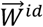 and 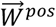 (Suppl. Fig. 2c).

To assess whether the recorded population was specialized, we computed the Spearman correlation between the two coding importance vectors, denoted as 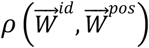. We then compared the value of 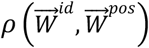 with those obtained by a null model where mixed selectivity is implemented by assigning two random coding directions in the neural space to the two variables. In this null model, the two coding vectors 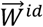 and 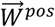 have no relationship with each other. To implement this null model, we sampled two random rotation matrices, one for each variable, and used them to rotate the weights of position and identity decoding before taking the average across cross-validations. This procedure was repeated 1000 times, each time with new rotation matrices, to obtain a population of null values (black curves in Suppl. Fig. 2d, f and black error bars in Suppl. Fig. 2 e, g). The recorded population of cells was then classified as either mixed or selective depending on the significance of the correlation, computed using the z-score of the Spearman correlation compared to the null model:

- 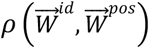 < null model: selective population (Suppl. Fig. 2a)
- 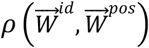 ~ null model: mixed selectivity (Suppl. Fig. 2b)
- 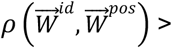 > null model: mixed selectivity (uneven distribution of coding importance, Suppl. Fig. 2c)

The whole procedure was performed for pseudo-simultaneous data (Suppl. Fig. 2 d, f) and for individual animals (Suppl. Fig. 2 e, g).

#### Cross-condition generalization performance

Cross-condition generalization performance (CCGP) was computed as described in ^16^. We first constructed pseudo-simultaneous activity vectors as described above, except we did not group data from pairs of conditions with the same decoding variable. Rather, pseudo-trials used for training a given classification came from one of the pairs of conditions that both contained the decoding dichotomy for a given classification while sharing the same non-decoding variable. The corresponding testing set consisted of data from the other pair of conditions that shared the non-decoding variable. For example, when decoding *social identity*, one training set consisted of data during interactions with mouse 1 versus mouse 2, when both were in the left cup, and the testing set consisted of data with mouse 1 and mouse 2 in the right cup. The decoding for a given dichotomy was then repeated, swapping the classes of pseudo-trials used for the training and testing data (e.g., training with data obtained with mouse 1 and mouse 2 in the right cup and testing on data with mouse 1 and mouse 2 in the left cup). CCGP was obtained from the mean decoding performance from the two pairs of training and testing conditions.

We estimated the null model CCGP as described in ^16^. To obtain a meaningful null model for generalization performance, it is important to maintain the level of decodability observed experimentally while selectively randomizing generalization between different pairs of conditions. To achieve this, we performed a solid rotation-translation of the pseudo-population vectors sampled from each condition in the neural activity space (using q=5 as described for the decoding analysis) by random shuffling of the neuron index. After the four independent rotations, we computed the CCGP as described above to obtain a null model CCGP value and repeated this to obtain 20 null model CCGP values. As described in the decoding section, the significance of the CCGP value for the experimental data was computed from its z-score with respect to the population of null model CCGP values.

#### Comparing decoding performance and CCGP across experiments

To compare the decoding performance or CCGP of the same subject in different experimental paradigms (for example, interacting with the two novel or the two familiar animals), we balanced the subject’s behavior so that each of the four conditions had the same interaction time (the minimum) between the two paradigms. If the two sessions had a different number of recorded neurons, say *n_min_* and *n_max_*, we randomly sub-sampled the session with a larger number of neurons to match the smaller one. The random choice of *n_min_* out *n_max_* neurons was repeated for each cross-validation (for decoding) or each pseudo-simultaneous data sampling (for CCGP) when decoding the *n_max_* session.

#### Null model for decoding difference

To assess the significance of a difference between two decoding performances and μ_*B*_, we first obtain a distribution of null model values for both performances as described above. We then created a null model distribution for the difference μ_*A*_ − μ_*B*_ by taking all possible differences between null model values for μ_*A*_ and null model values for μ_*B*_. The *p* value of the difference was then derived from the z-score of the performance difference μ_*A*_ − μ_*B*_ compared to this distribution of differences. Note that the null model distribution of differences has a standard deviation that is approximately the sum of the two standard deviations of individual null model distributions.

### Geometrical model

To test our geometrical interpretation of the experimental data, we developed a statistical model in which increasing degrees of familiarity led to a progressive and continuous change in the geometry of social/spatial representations. The model is composed of a population of *N* neurons whose firing rate is described by two binary latent variables, corresponding to position and stimulus identity of animals with the same degree of familiarity, reproducing the data from the interaction test with two novel or two familiar animals (Fig. 3).

In the absence of noise, each of the four conditions of an experiment would be associated with a point in N-dimensional neural firing space. To introduce response variability to the same stimulus, the population firing probability for each condition was described by an isotropic Gaussian distribution with unit variance centered around a condition-specific centroid in the neural firing space.

To account for our results during interactions with two novel animals, the means of the four gaussian distributions were arranged so that the two coding directions for the variables were orthogonal - reproducing a low-dimensional, or abstract, representational geometry in the firing space approximated by a two-dimensional rectangle. The length of two arms of the rectangle, denoted as 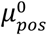 and 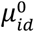, correspond to the signal-to-noise ratio in the representations of position and social identity variables, respectively, which in turn are reflected in decoding performance.

We accounted for the changes we observed in decoding of familiar compared to novel animals by introducing a *familiarity* latent variable, denoted as *f*, in which increasing degrees of familiarity modify the planar, rectangular representation of novel animals as follows.

1. Reduces signal-to-noise ratio of the *identity* variable: 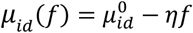
2. Performs a global shift by vector length *αf* along a third coding direction orthogonal to identity and position axes
3. Increases the representational dimensionality of the two variables by shifting each of the four condition centroids by a vector of length *γf* along a random direction for each condition

Using this model, we created simulated data for the activity of N neurons during a set of simulated sessions as a mouse is allowed to interact with two individuals of the same degree of familiarity, *f*, in left and right cups, with positions swapped in two trials. For each given condition (given mouse in a given cup), we randomly sampled T=5000 N-dimensional points from the distribution in neural activity space for that condition. We then analyzed the simulated data using the same linear decoding and CCGP procedures we used for the experimental data analysis. For each value of *f*, we repeated the sampling and analysis for *n* = 200 simulated sessions and took the mean and standard deviation for all decoding performance values (shown in Fig. 4e). We carried out this analysis for a set of values of *f* ranging from 0 (fully novel) to 1 (completely familiar) at increments of 0.1. We manually selected values of 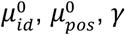, and *η* to reproduce the values of CCGP identity, CCGP position, and XOR decoding across the two-familiars and two-novels experiments at two optimized values of *f_NN_* and *f_FF_*, for a total of 6 fitted parameters to reproduce 10 experimental values: position decoding and CCGP, identity decoding and CCGP, and XOR decoding in the two-novels and two-familiars experiments. For the results shown in Figure 4, we used *N* = 80, 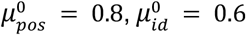, *η* = 0.45, *α* = 3.0, *γ* = 0.55, *f_NN_* = 0.23, and *f_FF_* = 0.78.

### Tradeoff between memory capacity and generalization in a Hopfield recurrent neural network

To study the trade-off between memory capacity and generalization capacity, we sampled patterns from geometries of different latent dimensionality, and measured (1) how many of these patterns a Hopfield recurrent neural network (RNN) can store and retrieve and (2) how well a linear classifier trained to decode a latent variable from these patterns is able to generalize across values of the other latent variables.

### Sampling patterns with varying dimensionality

To obtain patterns of varying dimensionality, we first defined *L* and *N* as the minimum and maximum dimensionality, with *N* ≫ *L*. A set of *P* binary *L*-dimensional latent patterns *λ^μ^,μ* ∈ [1,*P*] was then obtained by sampling *L* i.i.d. Bernoulli random variables *P* times:

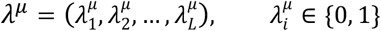

We then expanded each pattern *λ^μ^* into an *N*-dimensional embedding by repeating each 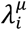 value *N/L* times, so that the collection of patterns kept the original dimensionality (*L*) in the new (*N*-dimensional) space.

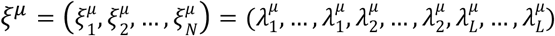

We call the L-dimensional space the *latent* space, with *λ_i_* being the i-th latent variable, and the N-dimensional space the *embedding* space, which corresponds to the neural activity space of N neurons.

We then increased the dimensionality of the patterns by randomly flipping each value 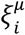 in each pattern with probability *δ*. Therefore, *δ* controls the dimensionality of the resulting set of patterns, which ranges from *L* (at *δ* = 0) to *N* (at *δ* = 0.5). We denote the collection of patterns obtained after applying this distortion as {*ξ^μ^*}”.

### Computing the memory capacity as a function of dimensionality

We then tested how *δ* affects the memory capacity of a Hopfield RNN of *N* neurons. Notably, when patterns are random and uncorrelated (in our case, 5=0.5) we expect a capacity that scales with the number of neurons *N*^38,39^. In Supplementary Information, we show through a theoretical argument that, in the case of patterns that span a *L*-dimensional space in an *N*-dimensional embedding (*δ*=0), the critical capacity is reached at O(L) patterns.

In the simulation shown in Fig. 1e, we numerically computed the memory capacity for the intermediate cases by varying 5 in 10 equally-spaced values from 0 to 0.5. For each value of *δ*, we constructed a set of *P* patterns {*ξ^μ^*}_*δ*_ as explained above. We then used these patterns to train a Hopfield model and tested its ability to retrieve each of the patterns used for training from a noisy version of the original (see Supplementary Information for more details). We then defined the maximum capacity of the model as the maximum value of *P* such that the fraction of retrieved patterns was larger than 95%. The resulting value of *P*(*δ*) for all 10 values of 5 was normalized with P(0.5) to be visualized in Fig. 1d.

### Computing the generalization capacity (CCGP) as a function of dimensionality

To compute how the generalization capacity of the neural code is affected by dimensionality, we used the same set of patterns {*ξ^μ^*}_*ξ*_ generated at a given value of *δ* and tested how well a decoder trained to report one of the latent dimensions generalizes across values of a second latent dimension *λ_m_*. For a given pair of latent dimensions (*l,m*), we divided the set of patterns {*ξ^μ^*}_*δ*_ into four classes depending on the value of the two latent variables. We then trained a linear SVM decoder to discriminate patterns in the (*λ_l_* = 0, *λ_m_* = 0) class from patterns in the (*λ_l_* = 1, *λ_m_* = 0) class, and tested it in the task of reporting patterns in the (*λ_l_* = 0, *λ_m_* = 1) class from patterns in the (*λ_l_* = 1, *λ_m_* = 1) class. This gave us a CCGP for the (*l,m*) pair, denoted as *CCGP_lm_*. We then took the mean CCGP value over all the possible pairs 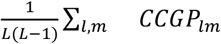 as a measure of generalization performance. This measure (ranging from a chance level of 0.5 to 1.0) was then normalized to range between 0 and 1 to be visualized in Fig. 1d.

## SUPPLEMENTARY FIGURES

**Supplementary Figure 1.**
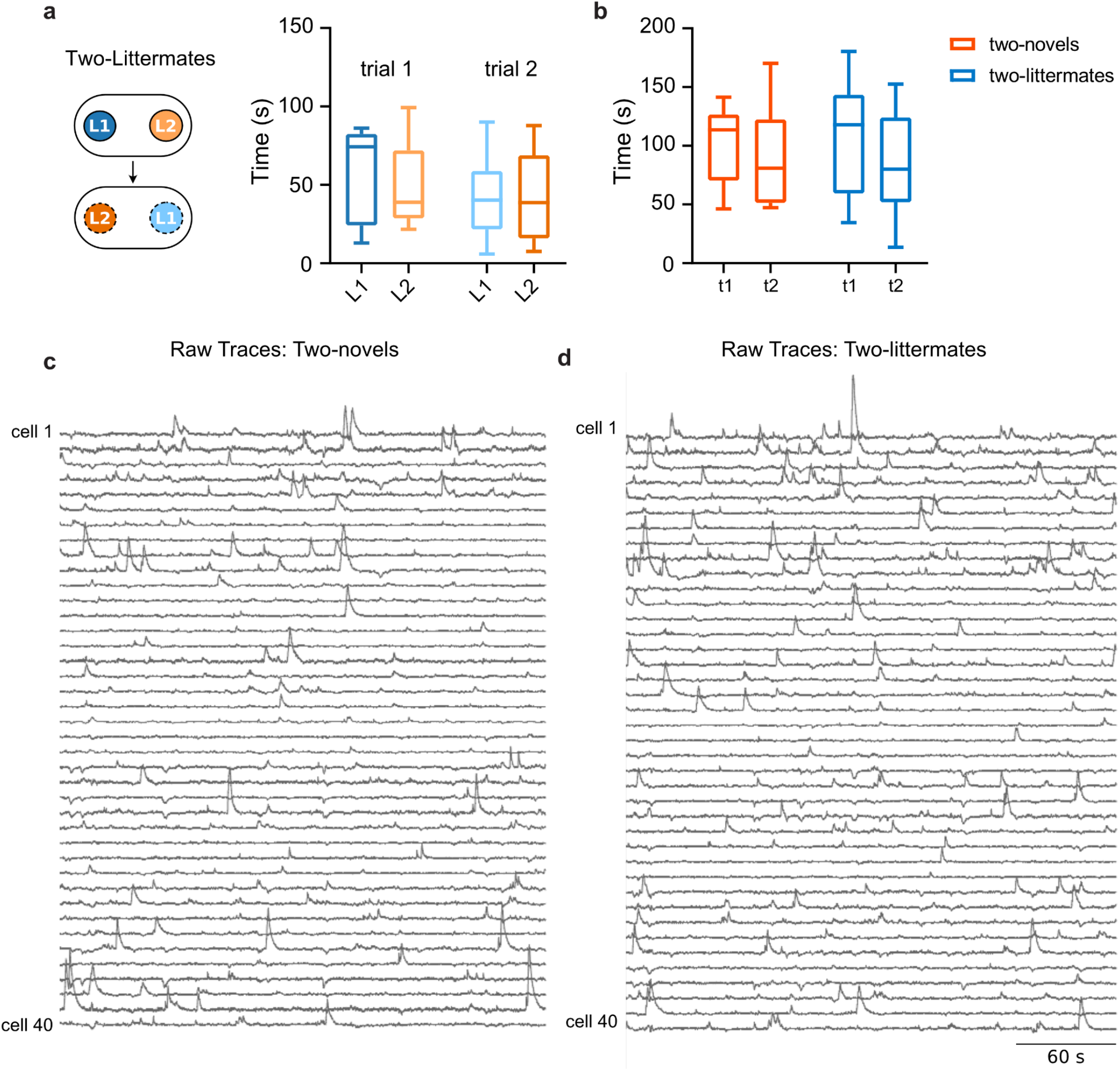
Behavior for the two-littermates experiment and comparison of behavior between the two-novels and two-littermates test. **a**) **Left**, experiment schema for two-littermates test. **Right**, total interaction time of subjects with the two littermate stimulus mice in the two trials. No significant difference was observed for either trial: Two-way ANOVA for Interaction Partner x Trial F(1,5)=0.2012, p=0.67. **b**) Comparison of summed interaction times with two novel mice (N1-N2) or two familiar littermates (L1-L2) in trials 1 and 2 for the experiments shown in Figure 1–3. No significant difference in interaction was observed between the two tests: two-way repeated-measures ANOVA of Test F(1,5)=4.342×10-5, p>0.99, or Test x Trial F(1,5)=0.5946, p=0.48. **c)** 40 example raw traces (dF/F normalized to trace maximum) from a single subject mouse across trial 1 of the two-novels test. **d)** 40 example raw traces from the same subject across trial 1 of the two-littermates test.

**Supplementary Figure 2.**
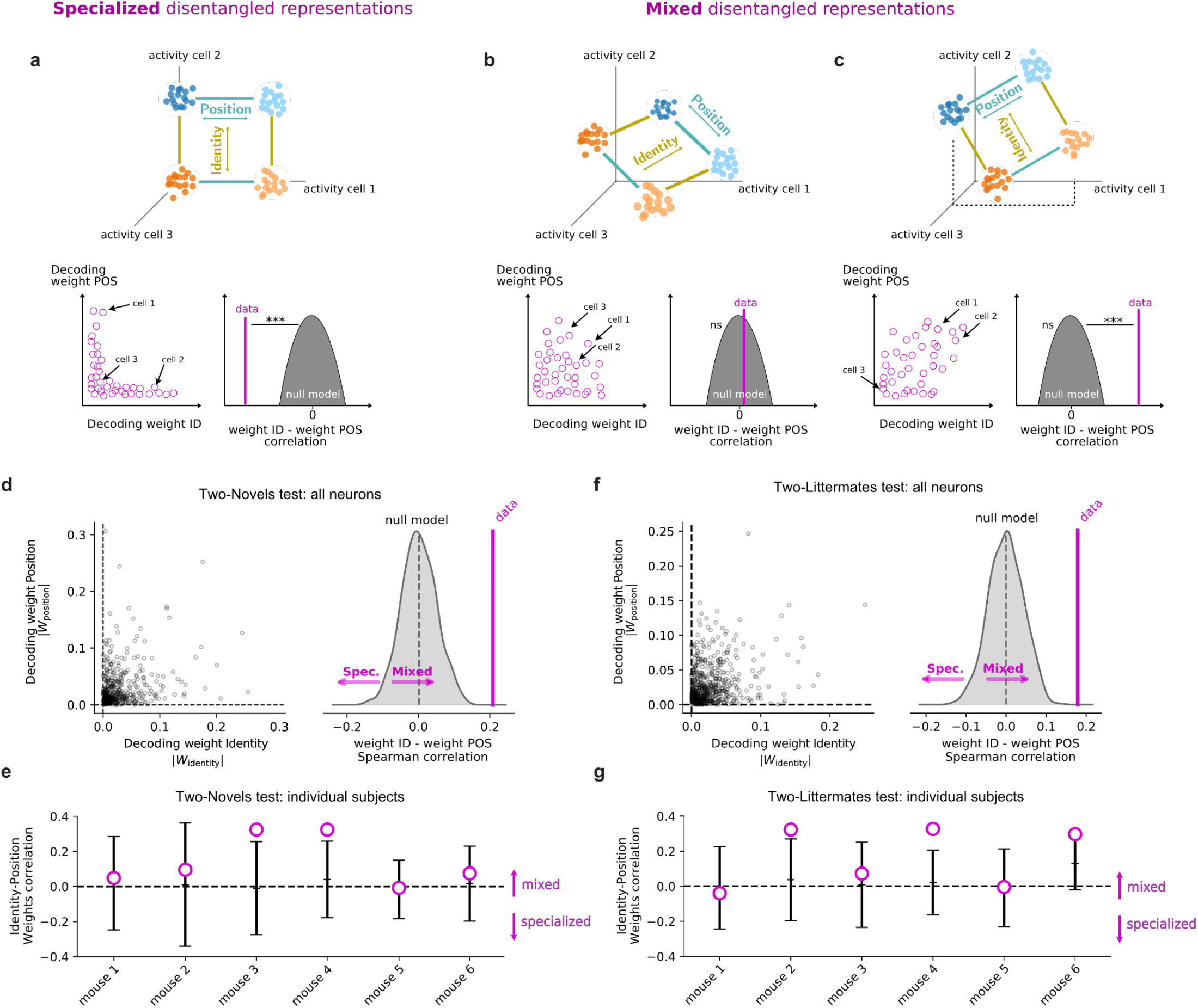
Decoding analysis indicates that CA2 neurons show primarily mixed selectivity for identity and position information. **a-c)** Three possible scenarios of neural selectivity for two variables: position (POS) and identity (ID) **a) Top**, geometric configuration in neural activity space for three theoretical neurons with selective firing, whose representation of the four conditions of our experiments is a low-dimensional plane with axes parallel to those of neurons 1 and 2. Neuron 1 responds selectively to position, neuron 2 to mouse identity, and neuron 3 to neither. **Bottom left**, schematic of decoding weights for POS versus ID for multiple hypothetical dCA2 neurons with selective firing, with positions of neurons 1-3 indicated as pictured in the top panel. In this scenario, we expect a significant negative correlation between the ID and POS weights compared to a null model where neural axes are rotated to enforce mixed selectivity (**bottom right**, see Methods). **b) Top,** geometric configurations for neurons with mixed selectivity to position and identity where information is mixed and evenly distributed across the population of cells. **Bottom**, in this scenario, there is no relationship between position and identity decoding weights (**bottom left**), therefore the ID-POS weights correlation is compatible with a null model (**bottom right**). **c) Top,** geometric configurations for neurons with mixed selectivity in which information is mixed but unevenly distributed across the population of cells (in the example, cell 3 does not encode position or identity information). **Bottom** In this scenario, there is a significant positive correlation between position and identity weights. **d) Left,** Experimental data plot for the pseudo-population of all subject mice of position decoding weights versus identity decoding weights from the test with two novel stimulus mice. **Right,** relationship of observed POS-ID weight correlation to a null model where neural axes are rotated to enforce mixed selectivity (see Methods). Results indicate that dCA2 neurons behave as in scenario **c**. **e)** Same analysis of panel **d** repeated for each individual subject mouse in the experiment with two novel mice. **f) Left**, experimental data plot for the pseudo-population of all subject mice of position decoding weights versus identity decoding weights from the test with two littermate stimulus mice. **Right,** relationship of observed POS-ID weight correlation to null model for pseudo-population of all subject mice. **g)** Same analysis of panel **f** repeated for each individual subject mouse in the experiment with two littermate mice. Null model error bars show mean ± 2 STDs.

**Supplementary Figure 3.**
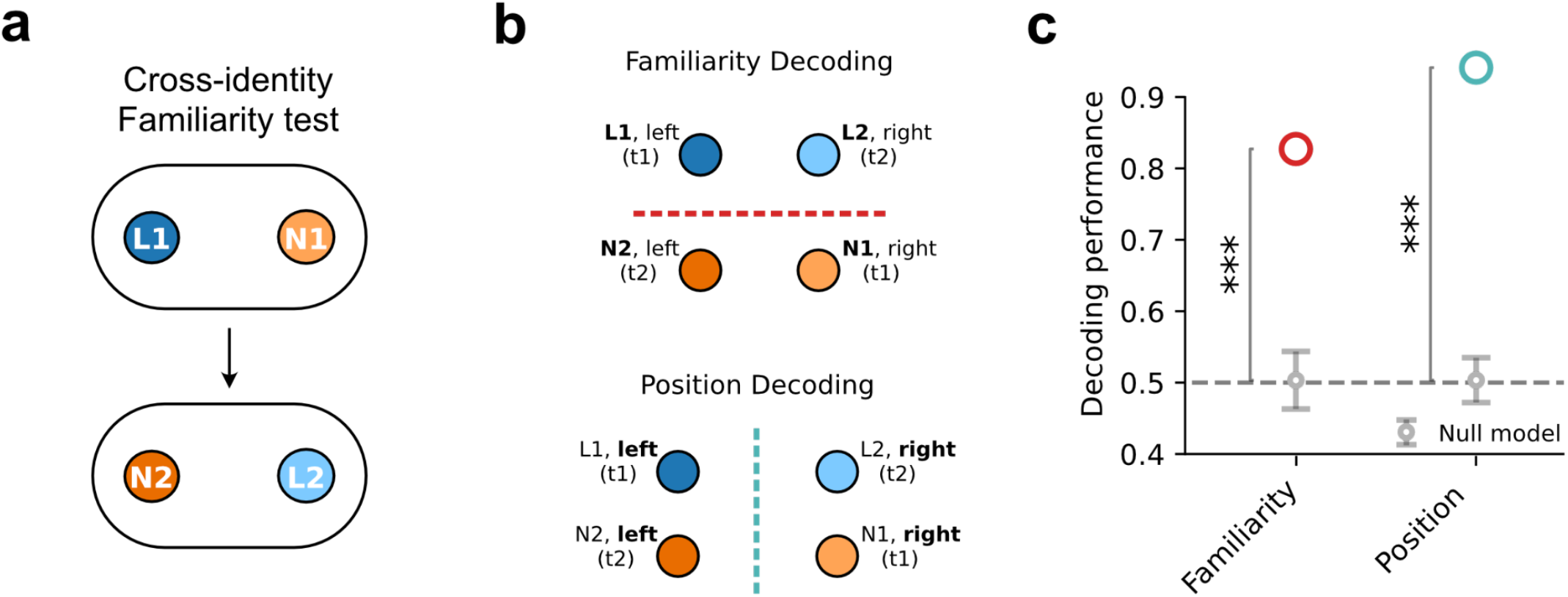
Decoding of social familiarity and position during experiment of Figure 4. n= 438 cells from 5 mice. **a)** Schema of experiment using two different pairs of novel animals and littermates in trials 1 and 2. **b)** Grouping of conditions for decoding social familiarity (top) and position (bottom). **c)** Performance of linear classifier for decoding social familiarity (familiarity decoding performance = 0.83, null = 0.50 ± 0.02 [mean ± SD, throughout figure], p<0.001) and position (position decoding performance = 0.94, null = 0.50 ± 0.02, p<0.001). Open circles show average decoding performance from 20 cross-validations. Gray dots and error bars show mean ± 2SD of distribution of chance values from shuffled data. P values are estimated from the z-score of the data values compared to the null model distributions.

**Supplementary Figure 4.**
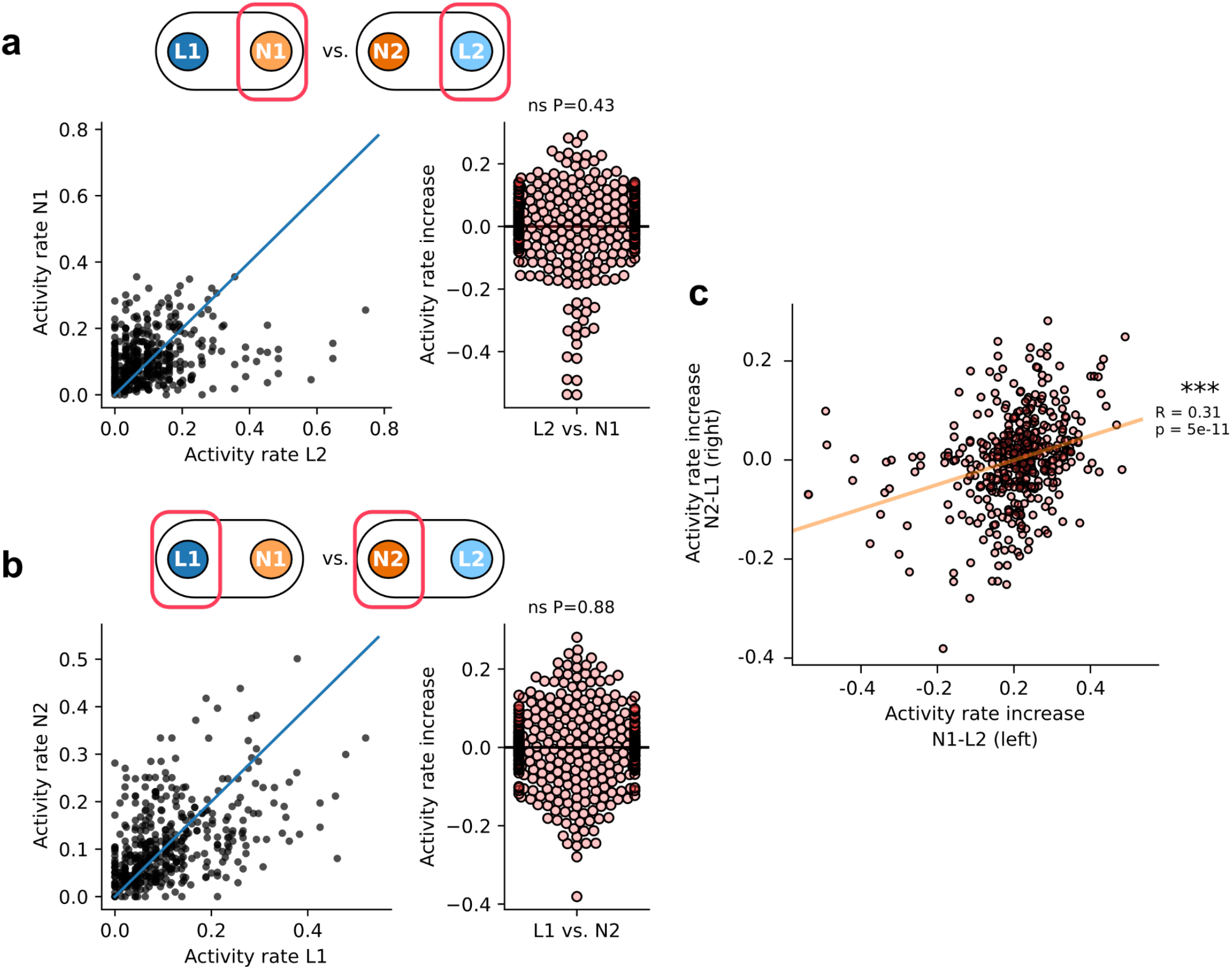
Mean activity rates, computed for each neuron as the fraction of 100 ms time bins with activity and rescaled to give a measure in Hz, of single neurons around novel versus littermate stimulus mice in experiment of Figure 4. **a) Left**, mean activity rate around the novel mouse N1 and littermate L2 across trials when both mice are located in the right cup. There is no significant increase in activity rate in the presence of the novel individual. **Right**, change in activity for each neuron across all animals around the novel mouse N1 and littermate L2 (mean activity rate increase from L2 to N1: −0.004 ± 0.11 Hz, one-way t-test, p=0.43). **b) Left,** Activity rate around the novel mouse N2 and littermate L1 across trials when both mice are located in the left cup. Again, there is no significant increase in activity rate in the presence of the novel individual. **Right,** change in activity for each neuron across all animals around the novel mouse N2 and littermate L1 (mean activity rate increase from L1 to N2: −0.0006 ± 0.09 Hz, one-way t-test, p=0.88). **c)** The difference in activity around N1 and L2 (both in the right cup) is correlated with the difference in activity around N2 and L1 (both in the left cup), consistent with an abstract representation of familiarity along a common random direction in the activity space (R=0.31, p=5×10^-11^). ns p>0.05; *** p < 0.001.

**Supplementary Figure 5.**
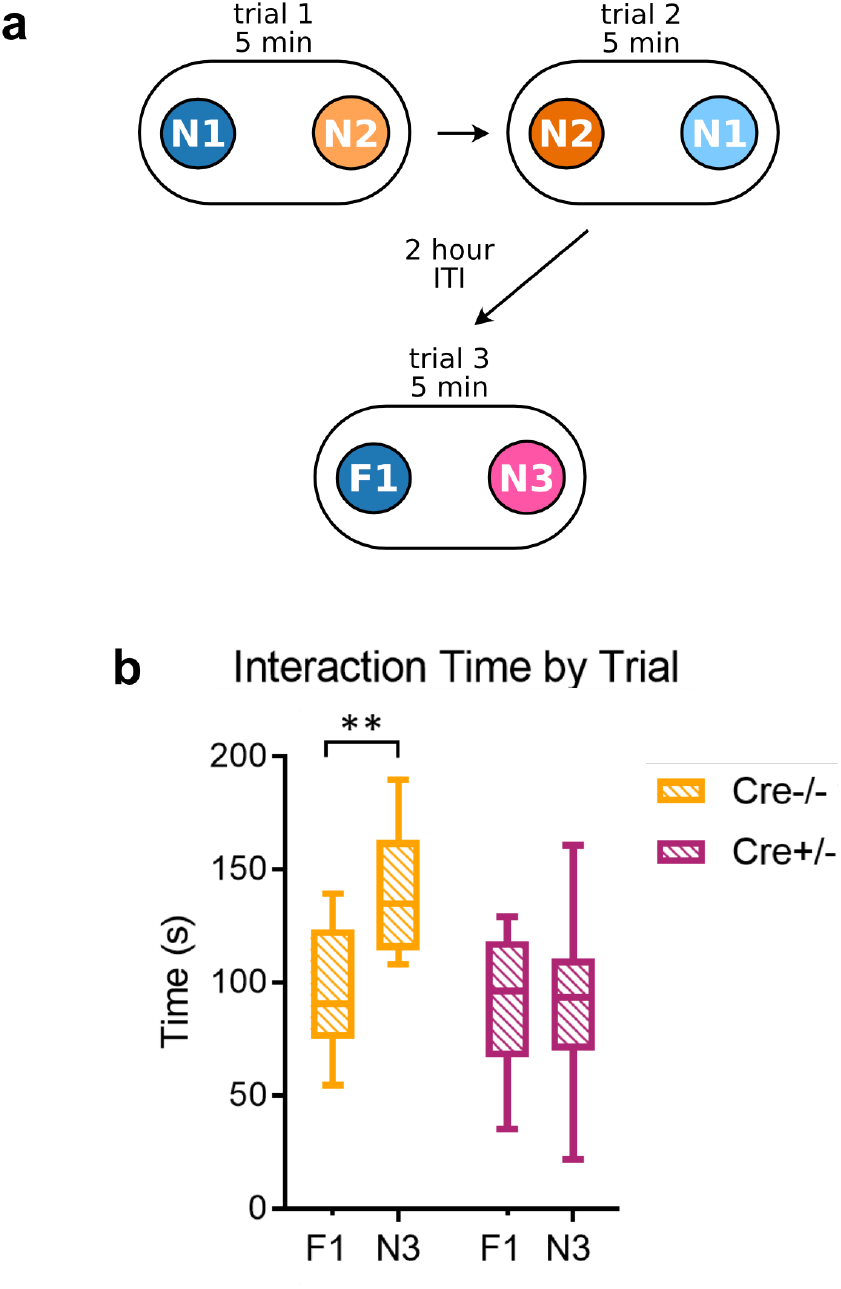
dCA2 silencing impairs social memory recognition in a two-choice test in an oval chamber. **a)** Experimental setup: Amigo2-Cre^-/-^ (control) and Amigo2-Cre^+/-^ mice were injected with Cre-dependent virus to express iDREADD in dCA2 in the latter subjects. Three weeks after viral injection both groups were systemically injected with CNO (5 mg/kg) 30-minutes prior to a social memory test, which consisted of two learning trials (trials 1 and 2) and one recall trial (trial 3). In trial 1, a subject mouse explored for 5 min two novel stimulus mice (N1 and N2) placed in pencil cup cages at opposite ends of an oval chamber. In trial 2, conducted immediately after trial 1, the positions of the two novel mice were swapped and the subject mouse explored the stimulus mice for an additional 5 min. Trial 3, After a two-hour intertrial interval, one of the two now familiar novel mice (eg N2) was exchanged for a third novel mouse (N3). Memory recall was assessed by the increased time spent exploring the third novel mouse compared to the now-familiar mouse from the previous trials (mouse N1, indicated as now-familiar mouse F1). **b)** Cre^-/-^ control mice showed expected increased interaction time with the novel compared to the familiar individual in trial 3. Cre^+/-^ mice, in which dCA2 was inhibited, did not show a preference for the novel over the familiar mouse. Two-way repeated-measures ANOVA: Genotype x Interaction Partner F(1,18)=6.435, p=0.021. Šídák’s multiple comparisons test. Trial 3 (F1 versus N3): Cre^-/-^ mice, p=0.0043, n=8; Cre^+/-^ mice, p=0.92, n=12. ** p<0.01.

**Supplementary Figure 6.**
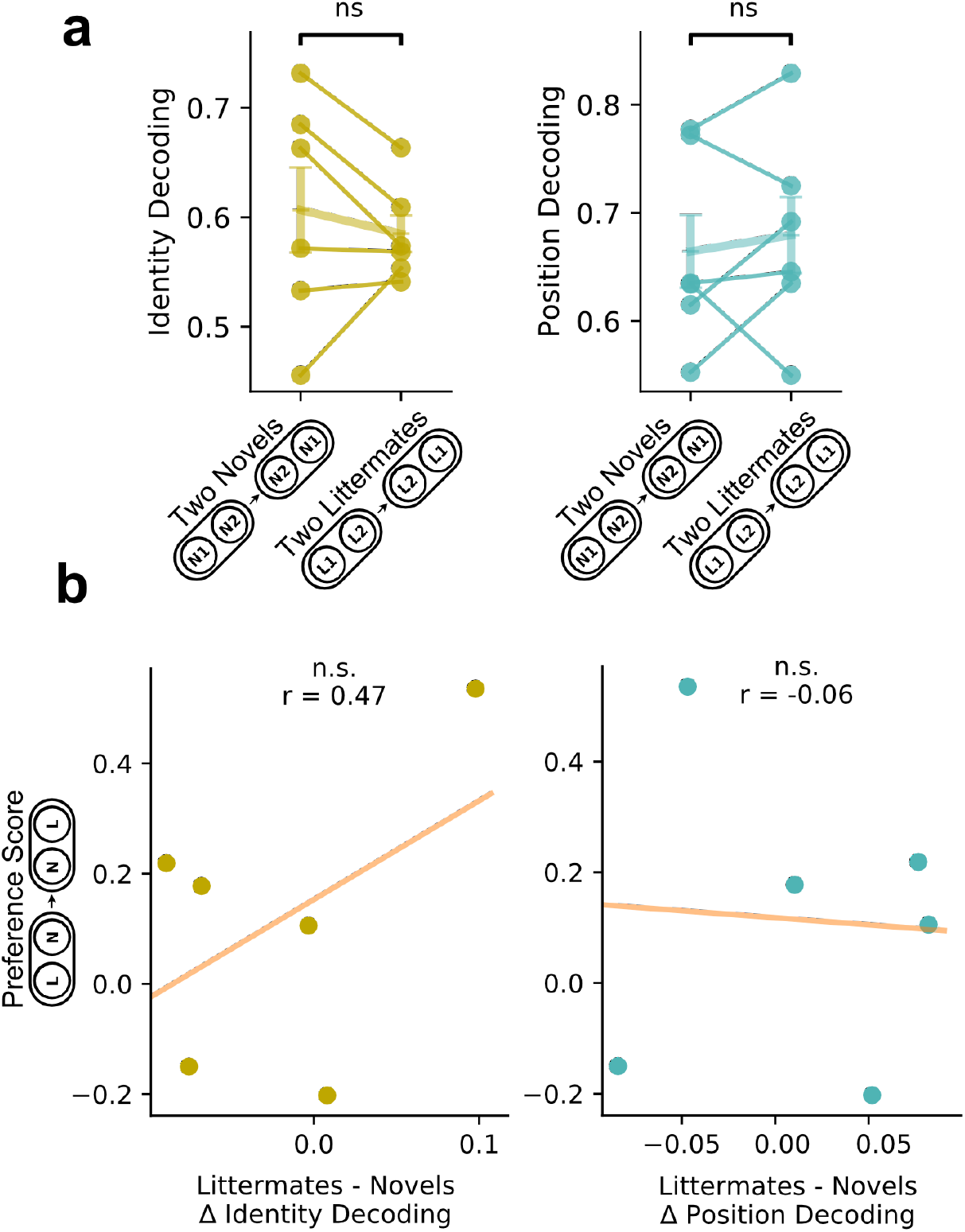
Comparison of identity and position decoding performance for individual subject mice in two-novel and two-littermate test with relation to behavioral preference in a social memory test. **a) Left,** there is no significant difference in identity decoding performance in tests with two novel (mean decoding performance = 0.61 ± 0.043 SEM) or two littermate stimulus mice (mean decoding performance = 0.59 ± 0.018 SEM; paired t-test p=0.49). **Right,** there is no significant difference in position decoding performance in tests with two novel (mean decoding performance = 0.66 ± 0.037 SEM) or two littermate stimulus mice (mean decoding performance = 0.68 ± 0.038 SEM; paired t-test p=0.61). Points represent individual subject decoding performance. Vertical and horizontal lines show mean ± SEM. **b)** There is no significant correlation between the difference in identity decoding performance (r=0.47, p=0.44, **Left**) or the difference in position decoding performance (r=-0.06, p=0.89, **Right**) in individual subject mice between the tests with two novel or two littermate stimulus mice and the preference score of the same subject mice for exploring a novel compared to familiar stimulus mouse conducted in a separate social memory test from Figure 5.

## Supplementary Information for “Distinct geometries of hippocampal CA2 representations enable abstract coding of social familiarity and social identity”

Our goal is to compare memory storage capacity of low- and high-dimensional representations. We assume that a memory of an experience is recollected when the neural circuit is presented with a cue and it can reconstruct the patterns of activity corresponding to the experience stored in memory. This can be implemented with a feed-forward network that essentially implements an autoencoder (see e.g. [1]) or in recurrent neural network like the Hopfield network [2, 3], in which each attractor of the neural dynamics represents one memory (this scenario would be compatible with the anatomy of dCA2, which is known to have recurrent excitatory connections [4]). In both cases, the synaptic weights are chosen in a way that the recollected memory is reconstructed: for the autoencoder the memory is simply reconstructed in the output layer, and for a recurrent network it is reconstructed after relaxation in an attractor. Also, in both cases a partial cue (e.g. a pattern that has a limited overlap with the one stored in memory) will lead to the reconstruction of the full stored memory.

In order to estimate the memory capacity we need to make assumptions about the nature of the memories. For random uncorrelated patterns the memory capacity of the Hopfield model is *p ~ N*: the number of attractors *p* scales linearly with the number *N* of neurons. Random patterns are high dimensional, as long as *p* is not too large (i.e. when *p < N*) and *N* is large enough, so this is one illustrative and highly representative case of memories that are represented with high dimensional geometries. Real world memories are not random and uncorrelated but it is not unreasonable to consider the random representations if one assume that the brain has a neural circuit that decorrelates the representations (recoding), at least so some extent, before storing them in memory (see for example [1]). This neural circuit could be implemented in the dentate gyrus, which is known to play an important role in pattern separation [5, 6, 7] (pattern separation is clearly a form of decorrelation).

### Theoretical derivation

We start by considering one possible way of constructing disentangled representations. The representations we now define are not the only possible type of disentangled representations, but they are a representative and illustrative example. Moreover, they have a geometry that is compatible with the observed low dimensional representations. Each pattern is obtained by concatenating *L* vectors of *N_L_* neurons, each encoding one latent variable Λ_λ_, with λ = 1, …,*L* (e.g. we could assume that *L* = 2 and the first *N_L_* neurons of the full vector encode the position of the animal, and the second *N_L_* neurons encode the identity). For simplicity we assume that each latent variable is encoded by the same number of neurons. All the neurons within each group of NL neurons have the same activation state, which equal to the value of the latent variable Λ_*λ*_ that they encode, and hence they are perfectly correlated. Following [2] we assume that there are only two activation states ±1 for each neuron.

The patterns to be memorized are 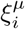 where *μ* is the memory index, *i* is the index of the neuron (*i* = 1, …,*N*). As discussed above, the patterns are obtained by concatenating vectors that encode different latent variables. Hence 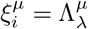 for *i* = (λ − 1)*N_L_* + 1,…, (λ − 1)*N_L_* + *N_L_*, where 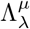 is value of the latent variable indexed by λ for memory *μ*. For example if *L* = 2, the memory *μ* would have the following form:

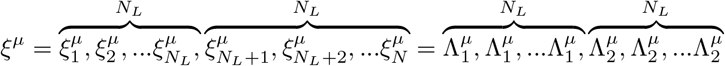

We assume that 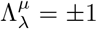 with equal probability. In other words the patterns 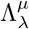 are random and uncorrelated. This implies that each memory is constructed by choosing randomly each latent variable. This could correspond to a particular episode in which, for example, a certain animal is encountered at a particular location. The identity of the animal and the location are assumed to be random. These representations are low dimensional as their dimensionality is *L* and *L* is assumed to be much smaller than *N*.

We now estimate the memory capacity using a simple signal to noise analysis, as in [2]. If the initial state is set by the input, and it is *s_l_* (*t*), then the state of activation at time *t* + 1 of neuron *s_k_* is given by the following expression:

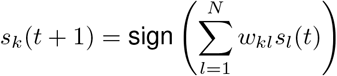

where *k, l* = 1, …*N* and *N* = *LN_L_* and *w_kL_* is the synaptic weight connecting neuron *l* to neuron *k*. The argument of the sign function is total synaptic current to neuron *k* and we call it *I_k_*. We assume that *w_kl_* is computed using the Hopfield prescription:

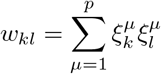

We now focus on the total incoming synaptic current to neuron *k*:

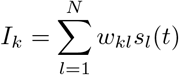

We consider the case in which a generic pattern is presented, for example memory 1: *s*(*t*) = *ξ*^1^. In the sum over *l*, we can now group together all the neurons that encode the same latent variable (they all have the same state of activation) and express the total synaptic current as a function of the Λ variables, which are independent by construction (both with respect to λ and to *μ*):

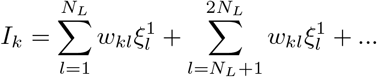

The first sum contains neurons that encode only the first latent variable Λ_1_, the second sum only the neurons that encode Λ_2_ etc. and all the states of activation 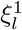 within each sum are the same: for example 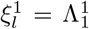 for all *l* = 1, …,*N_L_*. It is now convenient to switch to the indexes of the latent variables:

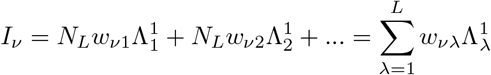

Where *v* is the index of the latent variable encoded by neuron *k*, the neuron whose state of activation has to be updated. *w_vλ_* is the value of the weight between a neuron encoding latent variable λ and a neuron encoding latent variable *v* and it is given by:

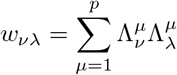

We then separate the sum over *μ* into two parts:

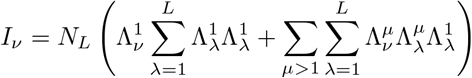

the first term reproduces the stored memory 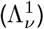 that has to be recollected and hence is usually called (memory) signal. The second accounts for the interference from the other memories, and under the assumption that the values of the latent variables are random and uncorrelated, it is basically just noise. As 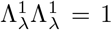, the signal scales like *N_L_L* and the noise term has a variance of approximately 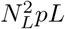 (there are *pL* independent terms in the noise). So the signal to noise ratio (SNR) is 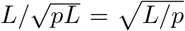. This means that the SNR of the memory to be recollected remains large enough, even in the presence of other memories, as long as *p* < *L*. Hence the maximum number of memories that can be recollected scales as *L*, the number of latent variables. Notice that *N_L_* cancels out, hence the max capacity *p* does not depend on the total number of neurons but only on the number of latent variables. This result is not surprising and it holds also for other learning rules. For example for the pseudo-inverse approach [8, 9, 3] it is clear that the memory capacity scales linearly with the dimensionality of the input patterns, which in our case is *L*.

### Numerical simulations

We verified this theoretical result by numerical simulations. Given a latent dimensionality *L* and a neural dimensionality *N*, we constructed p patterns Λ^μ^ by expanding the latent space into correlated chunks of *N_L_* neurons each in the neural space, as described above. We then used the *p* expanded patterns *ξ^μ^* to train an Hopfield model and tested its ability to retrieve one of the patterns used for training from a noisy version of the original. To construct noisy versions, we randomly flipped 10% of the units. A successful retrieval was identified if the model converged to a pattern with less than 5% flipped neurons compared to the original pattern after one step of the Hopfield dynamics, hence getting closer to the original pattern.

We then computed the fraction of retrieved patterns for different values of *L* and *N*. As shown in Figure 1a,b, the fraction of retrieved pattern decreases when p increases, a sign of limited memory capacity. The decreasing profile varied with *L* (Figure 1b) but not with *N* (Figure 1a), as expected by the theory. We then defined the maximum capacity as the maximum value of *p* such that the fraction of retrieved patterns was larger than 95% (green dashed line in Figure 1a, b). As shown in Figure 1c, this capacity increases with *N*, but it saturates to a value that depends on the latent dimensionality *N*. Moreover, this value is much lower than the one obtained with the same number of neurons in a high-dimensional setup where patterns are random and uncorrelated (black dashed line in Figure 1c). Finally, we computed how the maximum storage capacity, i.e. the maximum value of memorized patterns when *N* is large enough, scales with the latent dimensionality *L*. We found a good linear scaling of the maximum storage capacity with *L* (Figure 1d), hence confirming the results obtained in the theoretical derivation above.

### Limitations of the model

Notice that we had to assume that the weights between neurons encoding the same latent variable are all set to zero. Otherwise we have a problem similar to the presence of autapses in the Hopfield model (synapses that connect a neuron with itself): the autapses greatly enhance the stability of the input cue, at the expense of the ability to recall the stored memory [3, 9]. By setting all the synapses between neurons encoding the same latent variable to zero, we ensure that the network recollects the memory stored in the synaptic weights and it does not simply reproduce the cue. We neglected the corrections due to these zero weights in the formulae above because they do not change the scaling properties we are interested in when *L, N_L_* and *N* are large enough.

The simple calculations reported here have only the purpose to illustrate some properties of memory systems storing disentangled representations. It has several limitations: 1) the disentangled representations we considered are not the only possible low dimensional representations, and in particular we should consider representations that are rotated, which would be more similar to those observed in the experiment. In the simple case considered above each neuron encodes only one disentangled variable. 2) it will be interesting to consider representations that are not fully disentangled and have a dimensionality that is intermediate 3) the learning rule is very simple and it is biologically plausible but it doesn’t consider the problem of autapses (how does the system set to zero the connections between neurons representing the same latent variable?). On the other hand it seems to be clear from the experimental observations that CA2 is not really dealing with these low dimensional representations because the representations of familiar animals are high dimensional. The only purpose of the calculations reported here is to show that there is a problem of memory capacity with low dimensional representations and that is probably the reason why they are not used in CA2 to represent familiar animals.

**Figure 1:**
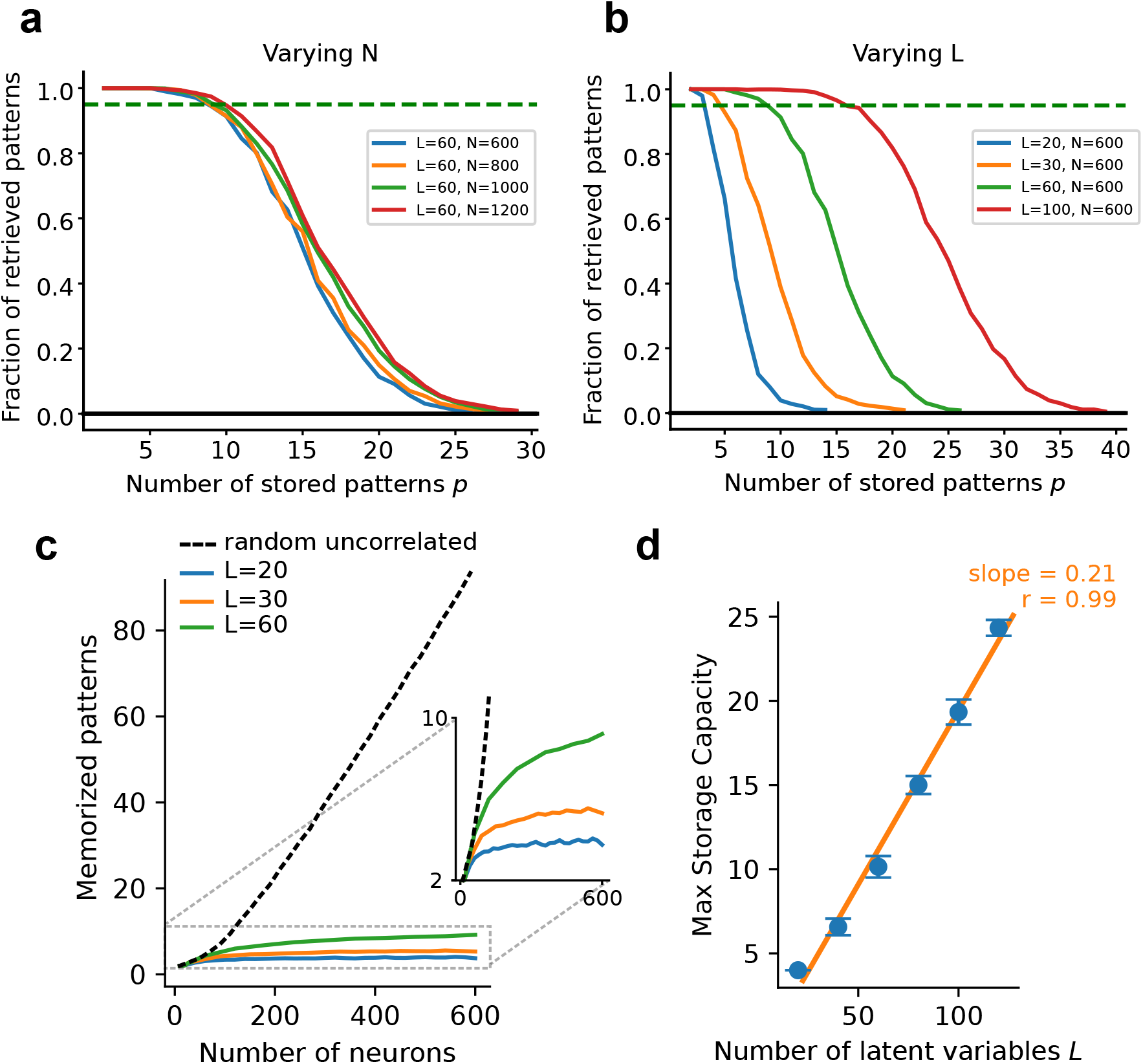
**a, b** Fraction of retrieved patterns as a function of the number *p* of patterns stored in the connectivity matrix of the Hopfield model. *L* denotes the effective dimensionality of the patterns, while *N* the embedding dimensionality (number of neurons). Curves obtained by averaging over *n* = 200 simulations for each value of *L, N*, and *p*. **c** Storage capacity, computed as the maximum number of pattern such that the model can retrieve at least 95% of them, as a function of the number of neurons N for a classical Hopfield model with random uncorrelated patterns (dashed black line), compared to the case of a low dimensional latent structure (*L* = 20 and *L* = 30). **d** max storage capacity, defined as the storage capacity of an Hopfield model of a large number of neurons (*N* = 2000) as a function of the latent space dimensionality *L*.

