## Supplementary figures and images for "Tuned geometries of hippocampal representations meet the demands of social memory"

### Supplemental Movie 2

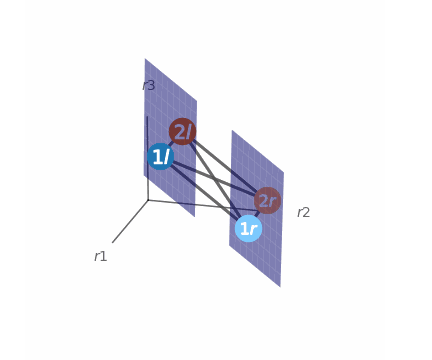

### Supplemental Movie 3

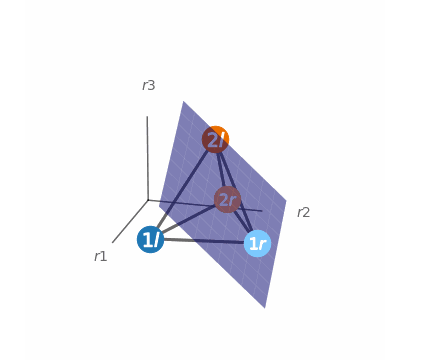

### Supplemental Movie 4

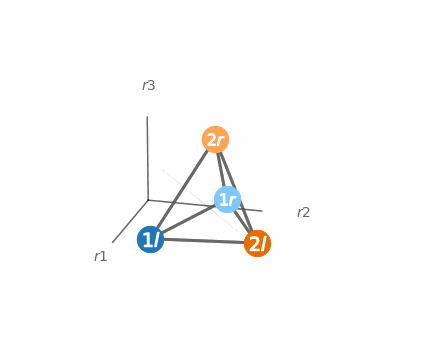
